# Comparing the impact of sample multiplexing approaches for single-cell RNA-sequencing on downstream analysis using cerebellar organoids

**DOI:** 10.1101/2024.08.23.609290

**Authors:** Kseniia Sarieva, Theresa Kagermeier, Vladislav Lysenkov, Zeynep Yentuer, Katharina Becker, Julia Matilainen, Nicolas Casadei, Simone Mayer

## Abstract

Sample multiplexing provides a solution to limited sample throughput in single-cell RNA sequencing (scRNA-seq) experiments. Different strategies for multiplexing are commercially provided by Parse Biosciences combinatorial barcoding (Parse) and 10x Genomics CellPlex combined with microfluidic cell capture (10x). However, the extent to which these two techniques differ when characterizing complex tissues such as regionalized neural organoids and whether data generated from the two techniques can be readily integrated is unknown. Cerebellar organoids are a highly relevant model for understanding evolutionary differences, developmental trajectories, and disease mechanisms of this brain region. However, they have not been extensively characterized through scRNA-seq. Therefore, we compared the two multiplexing techniques, 10x and Parse, using cerebellar organoids derived from three stem cell lines. While both strategies demonstrated technical reproducibility and revealed comparable cellular diversity including the main lineages of cerebellar neurons, we found more stressed cells in 10x than in Parse. Additionally, we observed differences in transcript capture, with Parse covering a higher gene biotype diversity and less mitochondrial and ribosomal protein coding transcripts. In summary, we demonstrate that both techniques provide similar insight into cerebellar organoid biology, but flexibility of experimental design, capture of long transcripts, and the level of cell stress caused by the workflow differ.

## Introduction

Single-cell RNA-sequencing (scRNA-seq) has revolutionized our approach to characterize cell types, states, and lineages in various biological systems and provides a new readout in screening applications and drug development^1,2^. Further, scRNA-seq is broadly applied to investigate cellular mechanisms in various model systems in health and disease^3^. The use of scRNA-seq has been limited by technically challenging workflows, often resulting in relatively low sample throughput in single experiments^4,5^. However, an adequate number of cells per sample and sufficient biological replica are essential for the success of single-cell transcriptomic studies. Effective cell sampling maximizes the capture of cellular heterogeneity, allowing for the precise identification and clustering of rare cell populations^6^. Large numbers of cells contribute to robust statistical power, facilitating the detection of subtle changes in gene expression. Biological replicates are crucial to distinguish true biological variability from technical noise, allowing reliable inference of cellular responses to experimental manipulations^6^. Recent advances in commercialized kits have overcome some of the technical obstacles limiting sample throughput by enabling sample multiplexing. Thereby both the number of cells assayed and the number of possible replicates or biological samples in a single experiment can be increased. The different approaches to multiplexing of scRNA-seq are characterized by differential sample throughput. Additionally, multiplexing can help detect multiplets and facilitates their removal prior to analysis^7^. While combinatorial barcoding is inherently multiplexed, microfluidic approaches require an additional labeling step for barcoding, mediated by antibodies or lipids^8^. Thus, from a technical perspective, multiplexing of samples allows upscaling experiments. However, increasing the number of samples remains technically challenging when working with fresh tissue because tissue dissociation, a highly manual process, needs to be parallelized^9^. Fixation of the dissociated cells before capture overcomes this obstacle, and different samples, for instance from different experimental time points, can be processed together, thereby avoiding batch effects of the capture.

Two commercialized approaches for sample multiplexing employ these different strategies and are commonly used by laboratories across the globe: 10x Genomics (hereafter, 10x) offers a microfluidic approach, while Parse Biosciences (hereafter, Parse) relies on combinatorial barcoding of fixed cells. The kits allow multiplexing of 12 (10x) or up to 96 samples (Parse). The higher the number of multiplexed samples, the lower are the per-sample costs of cell capture with both strategies. Despite their broad use in the scientific community, the two commercial technologies for multiplexed scRNA-seq, 10x and Parse, need to be compared in depth concerning their performance regarding differential transcript capture, cell type enrichment, and the amount of information that can be inferred from secondary analyses. A recent study compared both technologies using peripheral blood mononuclear cells (PBMCs) and demonstrated that Parse had a higher sensitivity for detecting rare cell types^10^. Furthermore, it was shown that Parse covered a wider range of gene lengths, and that 10x was biased towards more GC-rich transcripts^10^. However, it remains unclear, to what extent these differences apply and potentially affect downstream analysis of other cell types and highly complex tissue samples that require dissociation.

In parallel to scRNA-seq technologies, protocols for human induced pluripotent stem cell (iPSC)-derived organoids have been developed and have rapidly gained importance in biomedical research over the last decade^3,11,12^. Particularly, the establishment of brain, or neural, organoids has greatly impacted neuroscience research as they allow to investigate the developmental stages that usually happen *in utero* and are experimentally hardly accessible^13^. Over the last few years, the protocols for generating neural organoids have been modified to generate regionalized tissues resembling neocortex, midbrain and cerebellum^14–17^. Regionalized neural organoids are more homogeneous and contain specialized cell types compared to non-regionalized organoids and are, therefore, a particularly powerful tool to study human neurodevelopment^18^, to model neurological disorders^19,20^, and to test on– and off-target effects of pharmaceuticals^21,22^. Despite their advantages and broad application, they can be a challenging model system due to the heterogeneity between batches of differentiation and iPSC lines, the diversity of generated cell types, and off-target tissue^23,24^. These limitations highlight that neural organoids require comprehensive characterization of cell and tissue types at single cell resolution by high-throughput technologies such as scRNA-seq to exploit their full potential^3^. Further careful characterization of new protocols with multiple iPSC lines should be performed to ensure reproducibility across cell lines^25^. While neocortical organoids are broadly used and extensively characterized through scRNA-seq, much less data is available for other regionalized neural organoids such as cerebellar organoids^15,26–29^.

The human cerebellum has long been thought to mainly be involved in motor learning and coordination^30^, however more recent insights into cerebellar function, describe its major contribution to cognitive functions such as attention, task execution, working memory, language and social behavior^31^, and a role in neurodevelopmental disorders such as autism spectrum disorder (ASD)^32,33^. Considering that regionalized neural organoids, including cerebellar organoids, depict the cellular compositions and mechanisms of the developing human brain^3434343434343434^, they are a promising tool for studying neurodevelopmental disorders affecting the cerebellum. Early developmental stages of cerebellar ontogenesis are conserved across species, with two progenitor zones arising in the rhombencephalon r1^35,36^. These two progenitor zones are the ventricular zone (VZ) and the rhombic lip (RL). The VZ gives rise to all inhibitory neurons of the future cerebellum, including Purkinje cells (PC) and inhibitory neurons of the deep cerebellar nuclei. In contrast, the RL generates all excitatory neurons, including, for example, granule cells (GC) and excitatory neurons of the deep cerebellar nuclei^37^. However, the human cerebellum is uniquely characterized by features including changes in neuronal subtype ratios and folial complexity with respect to other mammals^37,38^. Moreover, a comparison between human and non-human primates has revealed the existence of a distinct basal progenitor population within the VZ and a longer persistence of neural progenitors originating from the RL in humans^39^.

To date, cerebellar disorders such as cerebellar hypoplasias, Dandy-Walker Syndrome, ataxias, and medulloblastoma have mainly been studied in mouse or zebrafish models^40–43^. Cerebellar organoid models now provide an interesting avenue to model these disorders in a human tissue context^28^, as pioneered in several recent studies^20,44,45^. However, the protocols underlying their generation are still being improved^15,27,29^, and few single-cell RNA datasets of selected cell lines are available^15,29,46^. Moreover, recent publications on cerebellar organoid differentiation performed scRNA-seq on only one iPSC line^27,29^. Hence the reproducibility for different iPSC lines is yet to be tested.

Taken together, (regionalized) neural organoids such as cerebellar organoids hold great potential to understand human-specific brain development in health and disease. However, these models can display heterogeneous results and efficiencies across batches and cell lines and require precise characterization of the cellular population and transcriptional profiles. Different scRNA-seq techniques have been reported to show individual strengths and weaknesses in PBMCs^10^. To investigate how these differences could potentially impact the analysis of scRNA-seq data of complex, heterogeneous and 3-dimensional (3D) tissue such as regionalized neural organoids, we generated cerebellar organoids from three iPSC lines and performed an in-depth comparison of 10x with Parse on both technical and biological levels.

## Results

### Experimental design and quality assessment

To assess the reproducibility of cerebellar organoid differentiation and comparability of two scRNA-seq methods, we differentiated three publicly available iPSC lines (BIONi010-C, BIONi037-A, and KOLF2.1J) into cerebellar organoids (Fig. 1a). All three cell lines were handled in parallel throughout the culture period. Samples were harvested at day 35 (D35) and day 50 (D50) of differentiation, and pools of 24 organoids per cell line and time point were dissociated into single cells. One aliquot of each cell suspension was used to perform 10x, the other to perform Parse scRNA-seq. For 10x, individual samples were labelled with cell multiplexing oligos (CMO), pooled, and then split into two lanes of a 10x chip and processed by 10x 3’ Gene Expression experimental pipeline (Fig. 1a). In parallel, the second aliquot was fixed according to the Parse protocol and stored at –80°C. Frozen samples of both time points (D35, D50) were subsequently subjected to combinatorial barcoding, and two sub-libraries were sequenced. For simplicity, we will refer to Parse sub-libraries as “libraries” throughout the manuscript. This experimental design enabled us to minimize the effect of biological variability and to focus on differences arising solely from the two techniques, 10x and Parse.

**Fig. 1.**
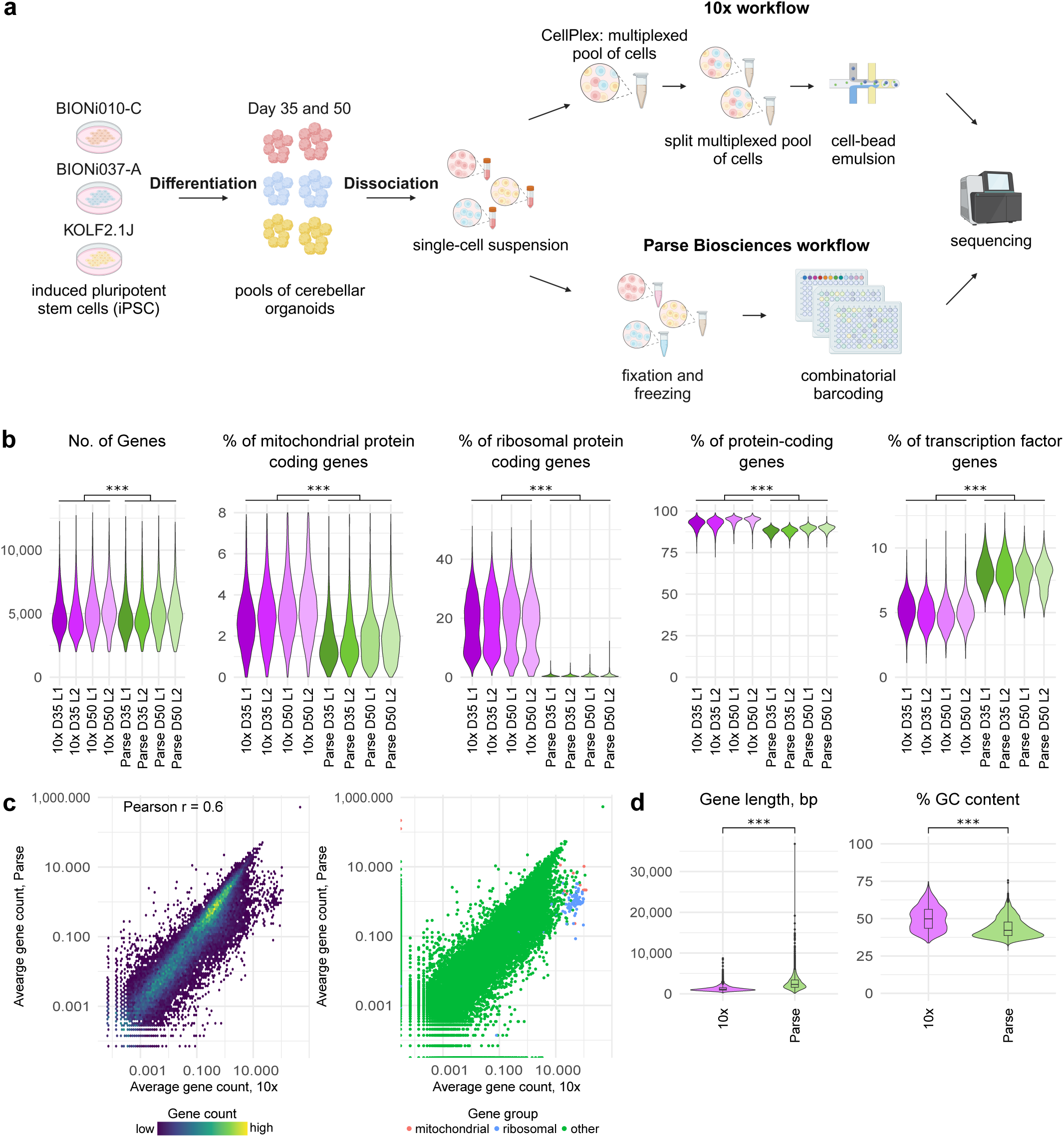
Study design, quality control, and potential biases in the data. **a**, Three iPSC lines (BIONi010-C, BIONi037-A, and KOLF2.1J) were differentiated to cerebellar organoids until days 35 and 50. The organoids generated from the same cell line were pooled and dissociated into single cells when each single-cell suspension was split into two portions. One set of single-cell suspensions was immediately subjected to sample multiplexing with CellPlex and processed in 10x Genomics 3’GEX+FB pipeline. The second set of single-cell suspensions was frozen until all samples were available. The samples were further processed though Parse Biosciences Evercode v2 pipeline. Libraries were sequenced, and the resulting FASTQ files were processed with technology-specific computational pipelines. Count matrices were further analyzed. Graphic was created with <u>BioRender.com</u>. **b**, Quality statistics after quality control. Color represents sample identity with respect to technology (10x or Parse), day of differentiation (D35 or D50), and library (L1 or L2). 10x, n = 29,505, Parse, n = 14,542 cells. Three-way ANOVA, p-values represent differences between technologies, *** p < 0.001. **c**, Left, density scatter plot showing correlation of average gene expression between the two technologies. Right, scatter plot showing correlation of average gene expression between the two technologies. Color represents gene group. **d**, Distributions of gene GC content and gene length for differentially expressed genes between technologies. Two-sided t-test, *** p < 0.001.

For both technologies, libraries were sequenced to achieve over 50,000 reads per cell (Supplementary Table 1). Due to varying sequencing depth, raw FASTQ files were downsampled to 50,000 reads per cell to allow a direct comparison of gene detection sensitivity (Supplementary Table 1). They were further processed through technology-specific alignment pipelines with human genome hg38: cellranger v7.2.0 multi pipeline for 10x samples and split-pipe v1.1.2 for Parse samples.

We first assessed the library efficiencies for both methods and found that, in both cases, most reads were mapped to the genome (93.2% for 10x, 91.8% for Parse, Supplementary Fig. 1a, Supplementary Table 2). While 56.3% of reads in 10x were mapped to exons, only 30.1% of reads were mapped to exons using Parse (Supplementary Fig. 1a, Supplementary Table 2). Valid barcodes were identified for 97.2% for 10x and 79.9% for Parse (Supplementary Fig. 1a, Supplementary Table 2). The cell recovery rate was 42.7% for 10x and 16.5% for Parse (Supplementary Fig. 1b, Supplementary Table 2).

To enable further comparisons, the technology-specific cell-by-gene count matrices were merged. We found that 32,408 genes had a non-zero expression in both technologies, while 2,159 and 12,098 genes were uniquely expressed in 10x and Parse datasets, respectively (Supplementary Fig. 1c). While the number of genes in the merged matrix was 62,910, it did not correspond to the number of genes with non-zero expression throughout the cells (Supplementary Fig. 1d). We therefore removed genes that had a non-zero expression in less than 8 cells in the merged count matrix. The resulting count matrix contained 38,580 genes.

For further analysis, we used the following combination of metadata parameters to assign cells to samples unless stated otherwise: (1) the technology (10x vs Parse); (2) the day of differentiation (D35 vs D50) of cerebellar organoids; and (3) the sequencing library (L1 and L2). Day of differentiation was used as covariate to acknowledge both biological differences in the stage of organoid differentiation and technical differences arising from harvesting D35 and D50 samples on different days. The sequencing library was used as a covariate to show the reproducibility of the workflow within each technology. In both technologies, libraries consisted of different cells, not different sequencing rounds.

Cell-level quality control (QC) was performed to remove cells with either a low or high number of detected genes, low number of genes per unique molecular identifier (UMI), and high percentage of mitochondrial protein-coding transcripts (Supplementary Fig. 1e). After QC, we recovered on average 87.2% of cells from 10x and 95.6% of cells from Parse datasets (10x, 29,505 out of 33,951 cells; Parse, 14,542 out of 15,226 cells). The number of detected reads per cell did not vary between the technologies before filtering because of the downsampling of reads approach we took to correct for differences in sequencing depth (p ≥ 0.05, Supplementary Fig. 1e). However, the number of genes per cell was higher in Parse both before and after QC (p < 0.001, Fig. 1b), suggesting that there might have been a higher diversity of detected gene biotypes in the Parse dataset. Indeed, while protein-coding genes were the most abundant gene biotype in both technologies, their percentage of the total reads was significantly smaller in Parse than in 10x (p < 0.001, 10x, 93.2 ± 2.9; Parse, 88.7 ± 2.1; mean ± SD) (Fig. 1b, Supplementary Fig. 1e). In contrast, Parse recovered a higher proportion of non-coding RNAs (ncRNA) reads, including long non-coding RNAs (lncRNA) (p < 0.001; 10x, 6.7 ± 3.1; Parse, 9.2 ± 2.3; mean ± SD) (Supplementary Fig. 1f), which have previously been shown to be informative for cell type identification^47^. The difference in ncRNAs and exonic reads can be explained by primers used for reverse transcription: Parse uses a mixture of random hexamer and poly-dT barcoded primers for reverse transcription^8^, while 10x uses only poly-dT primers.

Additionally, the percentage of mitochondrial and ribosomal protein-coding genes was lower in the Parse than in the 10x samples. In contrast, the percentage of reads originating from transcription factors (TF) among protein-coding genes was higher in the Parse than in the 10x dataset (Fig. 1b, Supplementary Fig. 1e). In line with the observation of higher gene biotype diversity in Parse data by Xie and colleagues ^10^, this suggested differential gene detection between the two technologies. Indeed, when we analyzed the correlation of gene expression between the two technologies across cells, we found only a moderate correlation, which corresponded to previous findings (Pearson’s r = 0.6) (Fig. 1c)^10^.

Different bulk and single-cell RNA-seq technologies are known to have biases in gene detection based on gene properties such as GC content and gene length^10,48^. To characterize these biases in our cerebellar organoid model, we analyzed the correlations between gene abundance and gene length or GC content, respectively, in both technologies (Fig. 1d, Supplementary Fig. 1f). When all expressed genes per technology were used for gene length and GC content analysis, small but statistically significant differences were observed (p < 0.001, Supplementary Fig. 1f). However, when we analyzed both parameters in differentially expressed genes (DEG) per technology (10x, 2,737 DEGs; Parse, 4,055 DEGs), we observed large differences in both gene length and GC content, reminiscent of previously published results (transcript length, bp: 10x, 1302.4 ± 728.0; Parse, 2715.9 ± 1754.8; GC content, %: 10x, 50.3 ± 8.2; Parse, 43.8 ± 6.5; mean ± SD)^10^. While a bias towards detecting longer genes in Parse was observed both for protein-coding genes and lncRNA, the difference was higher for the former (transcript length, bp: protein-coding genes, 10x, 1300.2 ± 720.3; Parse, 2901.1 ± 1748.9; lncRNA, 10x, 1352.9 ± 888.7; Parse, 1595.5 ± 1311.1; mean ± SD) (Supplementary Fig. 1g). Finally, we performed an extensive analysis of gene detection sensitivity and biases largely corroborating results from the previous benchmarking study on PBMCs^10^ (Supplementary Table 2) in a different sample type, human cerebellar organoids, therefore suggesting that the observed differences are characteristic features of 10x and Parse technologies independent of the sample type.

### Technical and biological differences between technologies

Following the scRNA-seq QC workflow described above, we normalized the data and revealed highly variable genes for further Principal Component Analysis (PCA) as well as Uniform Manifold Approximation and Projection (UMAP) on unintegrated data (Fig. 2a). As expected from previous results and our findings on the QC level, both PCA and UMAP revealed major differences between the technologies. We hypothesized that these differences may be arising from different sample preparation procedures between the technologies. Single cell suspensions for Parse analysis were immediately fixed and frozen after dissociation, while cells undergoing 10x capture were live cells depleted of nutrients from the media for longer (including time periods for multiplexing with CMOs, transportation to the sequencing facility, cell counting and viability assessment) and passed through microfluidic channels of the instrument before lysis.

**Fig. 2.**
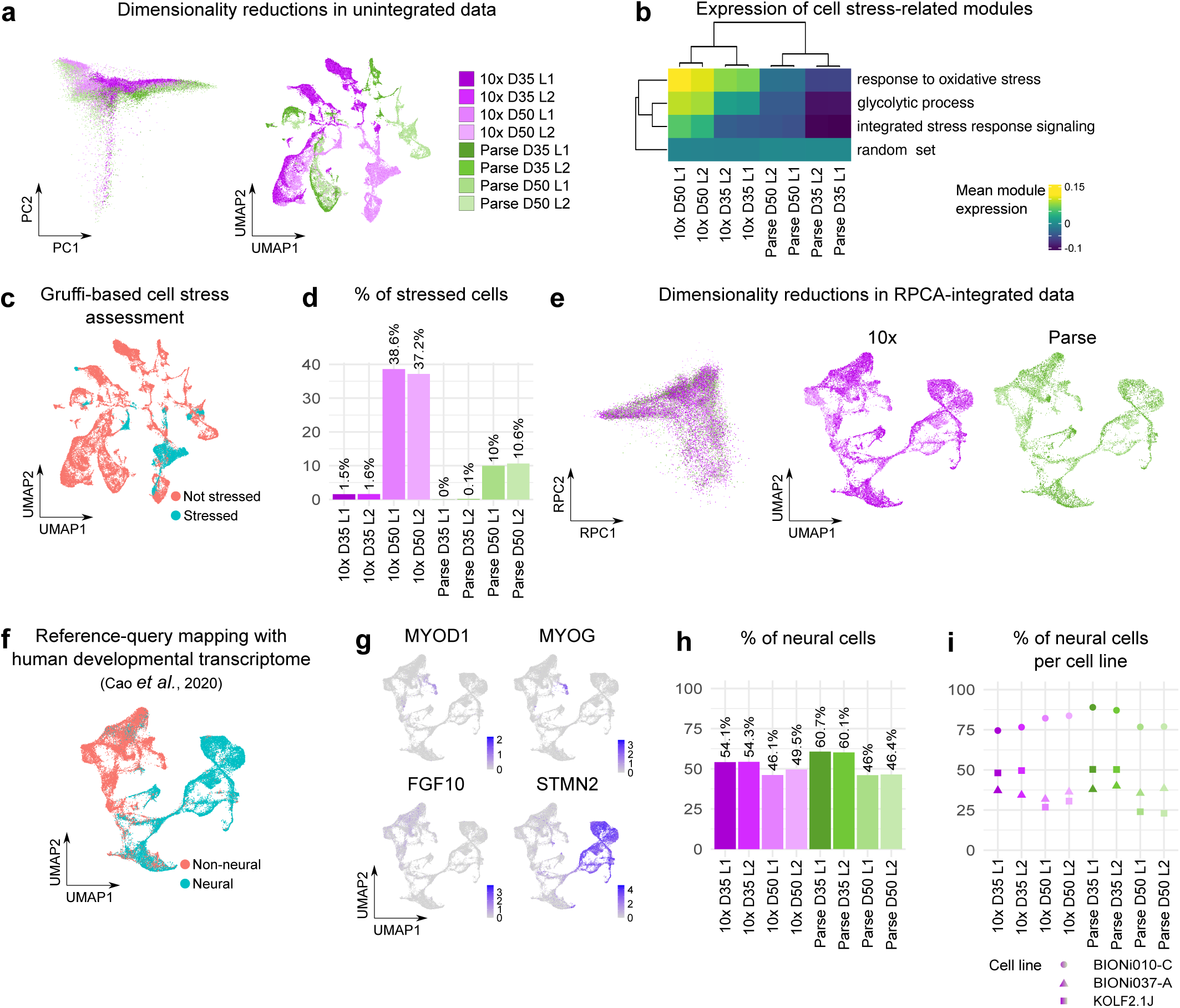
Assessment of neural lineage identity. **a**, PCA and UMAP plots for globally normalized and unintegrated data. **b**, Heatmap representing mean module expression scores of gene ontology terms related to aspects of cell stress. **c**, UMAP plot representing cell stress status of cells based on Gruffi assessment. **d**, Percentage of stressed cells based on Gruffi assessment. **e**, RPCA and UMAP plots for globally normalized and RPCA-integrated data originating from non-stressed cells. **f**, UMAP plot representing neural lineage status of cells based on reference-query integration with human developmental transcriptome^52^. **g**, Feature plots showing expression of selected genes to highlight developmental lineages. **h**, Percentage of neuroectodermal cells based on reference-query integration with human developmental transcriptome. **i**, Percentage of neuroectodermal cells per cell line based on reference-query integration with human developmental transcriptome. For **a, d, e, h, i**, color represents sample identity with respect to technology (10x or Parse), day of differentiation (D35 or D50), and library (L1 or L2).

Hence, we hypothesized that cellular stress may be a major contributor to differences between samples. We analyzed the expression of gene ontology (GO) modules involved in different modalities of cellular stress (e.g., GO terms for response to oxidative stress, cellular response to starvation) as well as downstream effects such as programmed cell death and integrated stress response (ISR) (Supplementary Fig. 2a). Module, or gene signature, expression analysis evaluates the expression of a set of genes rather than individual genes thereby providing hypothesis-driven insights into biological functions^49^. We included a random set of genes of average size of other gene sets into module expression analysis to serve as an internal control (Supplementary Fig. 2a). We performed hierarchical clustering of average GO module expression scores across samples, which revealed that samples from the two technologies clustered apart (Supplementary Fig. 2a). We also noticed that the major differences came from three cellular stress terms: response to oxidative stress, glycolytic process, and ISR signaling (Supplementary Fig. 2a). When using only these three modules and the random set for hierarchical clustering, the results were identical to the full list of cell stress terms (Fig. 2b, Supplementary Fig. 2a).

To understand the impact of cell stress on the dataset we further aimed to determine the number of cells with high cell stress transcriptomic signature. Therefore, we performed Gruffi cell stress assessment^50^ using two of the top cell stress terms from the module expression analysis: glycolytic process (GO:0006096) and ISR signaling (GO:0140467). With thresholds set to 95.5% quantile for GO:0006096, 89.8% quantile for GO:0140467 and no threshold for neurogenesis (GO:0022008) (Supplementary Fig. 2b), we found that the percentage of stressed cells varied between technologies but also between days of organoid differentiation (Fig. 2c,d). There were more stressed cells in the 10x data than in the Parse data, and both technologies captured more stressed cells in D50 than in D35 cerebellar organoids (Fig. 2d). This finding can be explained by the diffusion-based distribution of nutrients in organoids leading to an increasing nutrient deficiency as organoids grow bigger (D50 vs. D35)^23,51^. We therefore removed cells that were classified as stressed by Gruffi (6,595 out of 44,047 cells that passed QC) from further analysis, integrated normalized counts by sample using reciprocal PCA, and repeated PCA and UMAP. This analysis revealed that the data from the two technologies can be easily integrated (Fig. 2e).

To analyze the biological reproducibility of the cerebellar organoid protocol between different iPSC lines, we characterized the cellular diversity within organoids. We first aimed to understand whether organoids had neural identity. We therefore performed reference-query mapping of our dataset onto the human developmental transcriptome using Azimuth^52,53^. The reference dataset contained cells from 15 organs of human fetuses at 72 to 129 days post-conception, and the cells were captured using sci-RNA-seq3^52^. We first assigned our cells with cell types from this dataset^52^ (Supplementary Fig. 2c). The mapping score was high (0.71 ± 0.17, mean ± SD) (Supplementary Fig. 2d), indicating that our dataset corresponded well to the reference dataset^53^. However, the prediction scores varied between cells (0.59 ± 0.26, mean ± SD), with most cells not reaching a high-confidence prediction score of 0.75^53^. Given the relatively low prediction scores, we did not rely on specific annotation to certain cell types but further grouped the cells into two categories – neural and non-neural (Fig. 2f, Supplementary Table 3). We found a considerable portion of cells having non-neural identity (Fig. 2f) with subsets of cells expressing muscular markers (e.g., *MYOD1* and *MYOG*^54^) and endo-/mesodermal markers (e.g., *FGF10*, mesenchymal marker^55^) (Fig. 2g). In contrast, most cells classified as neural expressed the pan-neuronal marker *STMN2* (Fig. 2g). Overall, the proportion of neural cells ranged from 46.0% to 60.7% per sample (Fig. 2h). Importantly, considerable differences were observed between the three iPSC lines that the organoids were generated from (Fig. 2i). The BIONi010-C cell line had the highest number of neural cells (range, 74.5 to 89.0%), while KOLF2.1J-derived cerebellar organoids had 23.0 to 50.3% neural cells (Fig. 2i). Interestingly, D35 KOLF2.1J samples had about 50% neural cells, while at D50 only about 25% of cells were identified as neural (Fig. 2i) indicating that cells with neural identity do not proliferate further or die in comparison to other lineages.

To cross-validate our assignment to neural and non-neural cells, we adapted Gruffi^50^ for detecting neural and non-neural transcriptomic signatures. For that, we used GO terms for endoderm (GO:0001706, 57.8% quantile threshold) and mesoderm (GO:0001707, 66.7% quantile threshold) formation for selecting non-neural cells and GO terms for nervous system development (GO:0007399, 65.7% quantile threshold) and neurogenesis (GO:0022008, 64.8% quantile threshold) for selecting neural cells (Supplementary Fig. 2e). The results between reference-query mapping and Gruffi were coherent (82.6% classification overlap, Supplementary Fig. 2f). Inconsistent annotations between the two approaches were observed for putatively muscular cells (positive for *MYOG* and *MYOD1*) which were incorrectly classified as neural by Gruffi. We suggest that this discrepancy may be due to the shared excitability between neural and muscular cells.

### Characterization of neural cell diversity

Based on the reference-query mapping with the human developmental transcriptome^52^, we subset neural cells (19,526 neural cells out of 37,452 cells) and additionally downsampled 10x and Parse datasets to an equal number of cells (resulting in 7,212 cells per technology). We subsequently performed the integration and dimensionality reduction approach as described above.

Following developmental patterning *in vivo*, various experimental setups *in vitro* have found that forebrain structures develop upon neural induction, unless exposed to caudalizing factors^56^. Additionally, the gene expression program for telencephalon regionalization was upregulated in the cerebellar organoid protocol we used in the current study^26^. Hence, we aimed to reveal the brain regional identity of the neural cells. We analyzed the correlation of brain regional marker gene expression between our dataset and human brain transcriptomic data from postconceptional week (PCW) 12-13 from BrainSpan^57,58^. We used the list of brain regional markers compiled from the top 10 markers of different brain regions based on gene expression in E15 mouse brain^57^ (Supplementary Table 4). We found that all our samples had the highest correlation with the cerebellum (Supplementary Fig. 3a). However, when similarity scores were not scaled, we noticed that they were higher for 10x than for Parse samples (Fig. 3a). Next, we aimed to assign cell type identities to the neural cells. Combining cerebellar canonical marker gene expression^37–39,59^ combined with differential gene expression (DGE), we identified both RL-derived cellular lineages (RL, granule precursor cells (GPC), and GC) and VZ-derived newborn PCs (Fig. 3b,c). A subset of neuronal cells was characterized as hindbrain neurons, and we were not able to further refine our annotations (Fig. 3b). While overall proportions of cells captured by the two technologies were similar (Fig. 3d, Supplementary Fig. 3b), dividing progenitors, PAX6-positive RL and dividing RL cell populations were significantly enriched in the Parse dataset. In contrast, 10x captured more cells in a population that we could not annotate (Unknown 2) (Supplementary Fig. 3b).

**Fig. 3.**
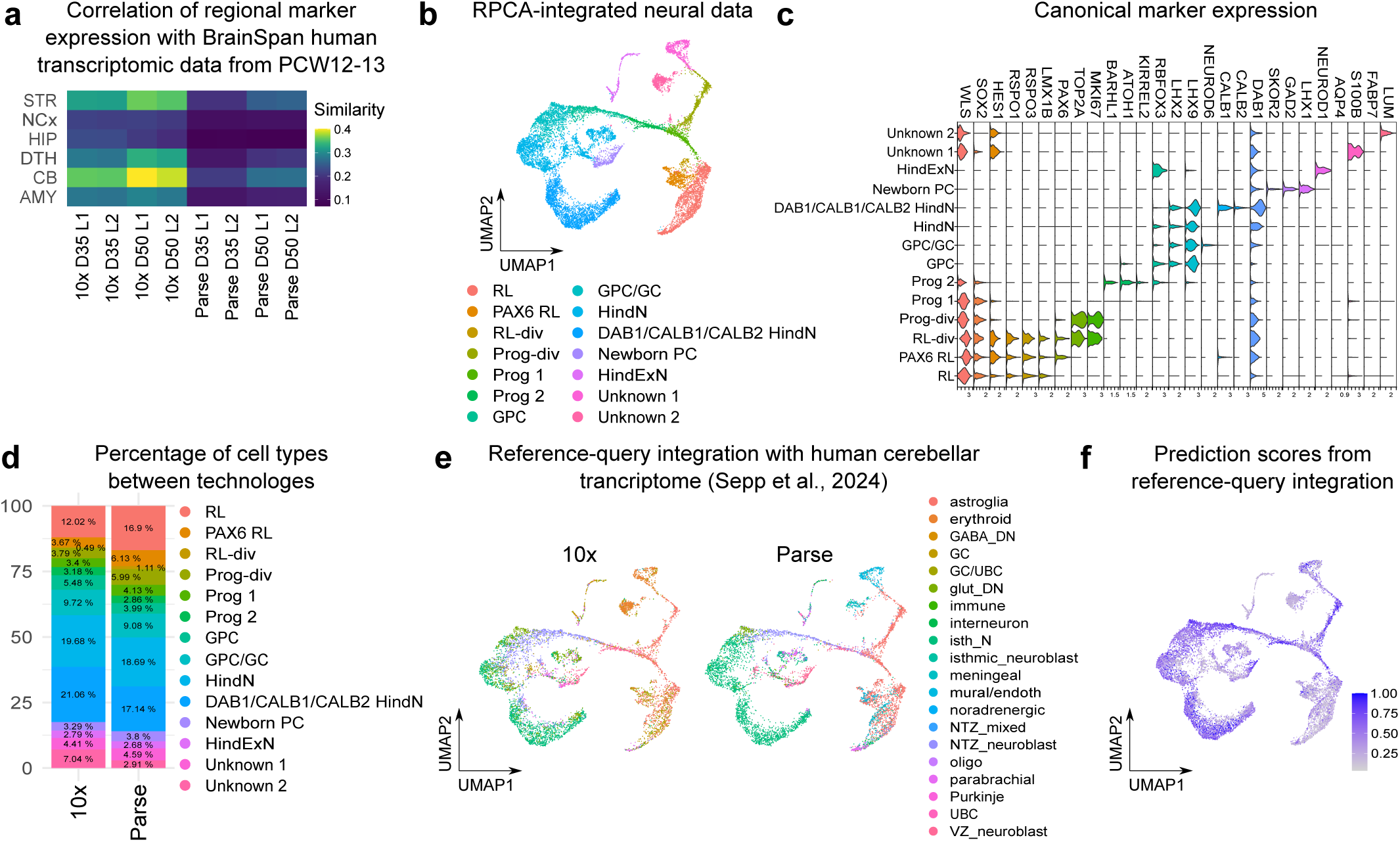
Assessment of regional identity and cell type annotation. **a**, Heatmap of similarity metric of VoxHunt algorithm comparing samples with human neocortical RNA-seq data from BrainSpan using brain regional markers obtained from Mouse Brain Atlas at E13. **b**, UMAP plots for globally normalized and RPCA-integrated neural data with manually annotated clusters. **c**, Violin plots for expression of canonical markers of hindbrain development. **d**, Stacked bar plot representing average proportion of individual cell types between technologies. **e**, UMAP plot representing cell type identity as assigned based on reference-query integration with human cerebellar transcriptome^38^. **f**, Feature plots showing prediction score based on reference-query integration with human cerebellar transcriptome.

To characterize the similarity of our cerebellar organoids with the developing human cerebellum, we performed reference-query mapping with the cerebellar transcriptomic dataset generated by Sepp and colleagues ^38^. To ensure that we compared our organoid data with early developmental stages of human cerebellum, we subset the reference dataset^38^ to only include prenatal samples. While finding a general agreement in cell type annotations, we noticed some differences both in assigned cell type identities (Fig. 3e) and prediction scores, which were higher in Parse than in 10x data (Supplementary Fig. 3c). One example of a discrepancy in assigned cell identities was RL cells of different subtypes. These cells were annotated as a plethora of cell types of the human cerebellum (Fig. 3e) and differed between 10x and Parse (Fig. 3e) but with very low prediction scores (Fig. 3f). We believe that the cause for this discrepancy may be that the reference dataset does not have a separate cluster for RL cells^38^. Instead, RL cells are part of an astroglia cluster consisting of both astroglia and RL cells^38^. The first separate cluster for RL-derived lineage was nuclear transitory zone neuroblasts (NTZ neuroblast)^38^, and in our dataset, the cells annotated as NTZ neuroblasts belonged mostly to progenitor 1 and GPC and GPC/GC clusters (Fig. 3b).

We further compared our data with the transcriptomic profiles of organoids from the recently published cerebellar organoid differentiation protocol (Supplementary Fig. 3d,e)^29^. The prediction scores were overall higher than for the comparison with the human cerebellar developmental transcriptome (Supplementary Fig. 3e). This time, however, prediction scores were higher for 10x than for Parse cells (Supplementary Fig. 3f). Interestingly, both reference datasets were obtained from the 10x pipeline, so the discrepancy in prediction scores between our Parse and 10x cells cannot simply be attributed to different technologies used for the generation of reference datasets. Instead, expectedly, our organoid data aligns more with organoid data obtained from a different protocol, than with primary tissue.

In summary, we found that the cerebellar organoids indeed acquired a mid-gestational human cerebellar regional identity. We also found robust differentiation into both major cerebellar lineages, RL– and VZ-derived cells. Small differences in the different parameters were found between 10x and Parse technologies.

### Secondary analysis between techniques reveals differences in cell stress signatures and neurodevelopment-related gene regulatory networks activity

In our QC, we found differences in the percentage of reads originating from ribosomal and mitochondrial protein-coding genes between the two technologies (Fig. 1b). We also found a subset of cells expressing cell stress-related genes, and the proportion of these cells was higher for 10x cells (Fig. 2d). Therefore, we next aimed to analyze whether the neural cells preserved these transcriptomic features and performed DGE analysis between the different technologies within individual cell types. For that, we split the dataset by cell type, technology, cell line, and day of differentiation and pseudobulked them for DESeq2. Overall, we found DEGs across all cell types (Fig. 4a, Supplementary Fig. 4a). Especially mitochondrial and ribosomal protein-coding genes were upregulated in 10x compared to Parse across cell types (Supplementary Table 5), including GPCs (Fig. 4b). More genes were upregulated in 10x compared to Parse across all cell types, further highlighting that with equal sequencing depth, 10x captures a lower variety of genes with larger numbers of reads per gene. Interestingly, there were a few genes with large fold change and relatively large p-values upregulated in either of the two technologies. These genes were identified as expressed either in 10x or Parse, as revealed by removing these genes from volcano plots (Supplementary Fig. 4b). To functionally characterize the differences in gene expression between the techniques, we performed gene set enrichment analysis and clustered the output by semantic similarity matrix (Fig. 4c). Here we describe findings for gene set enrichment analysis in GPCs, as a representative cell type with relatively high cell numbers and a medium number of DEGs. In GPCs, the normalized expression score for all statistically significant GO terms was less than 0, indicating their upregulation in 10x compared with the Parse dataset (Supplementary Table 6). Among these GO terms, we found a cluster of enriched GO terms related to nucleotide processing as well as a cluster related to mitochondrial respiration. These two clusters of GO terms included not only mitochondrial protein-coding genes as defined in scRNA-seq quality control (i.e., starting with “MT-”, Fig. 1b) but also other genes involved in mitochondrial function, for example, the *NDUF* gene family, which encodes nuclear-encoded genes coding NADH dehydrogenase (ubiquinone) subunits. Another group of enriched GO terms in GPCs was described as related to neuron projection assembly (Fig. 4c).

**Fig. 4.**
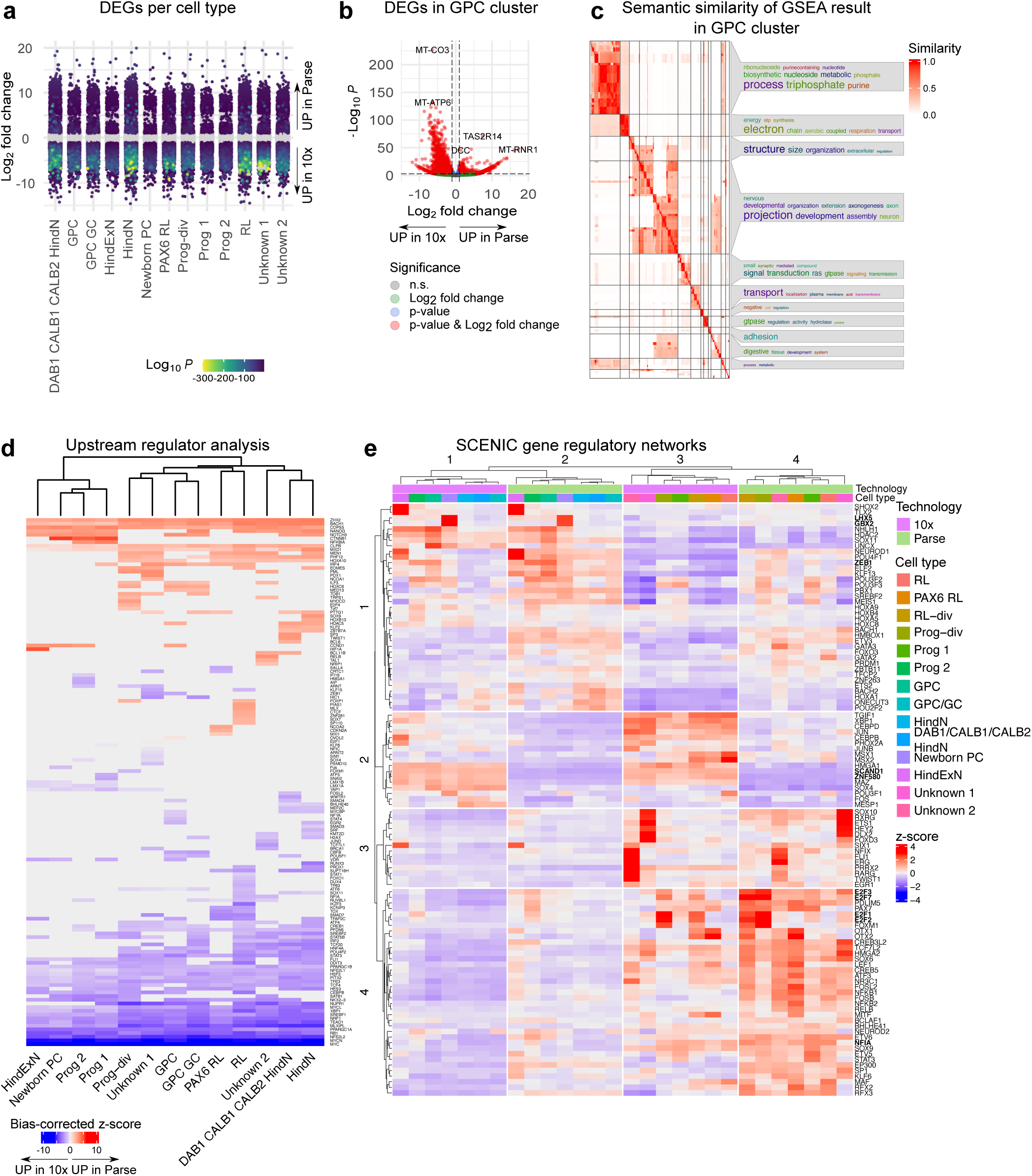
Differential gene expression between technologies. **a**, Strip plot displaying DEGs between technologies per cell type. Genes represented in grey are not differentially expressed. Color represents log10 adjusted p-value for differentially expressed genes (absolute log2 fold change > 1, FDR < 10^-4^). **b**, Volcano plot representing differential gene expression in GPC cluster. **c**, Heatmap representing semantic similarity between GO terms identified as significantly deregulated in GPC cluster by GSEA analysis. **d**, Heatmap representing z-scores for SCENIC regulon activity calculated based on AUC scores.

To reveal the upstream mechanisms leading to the described transcriptional changes across cell types we used ingenuity pathway analysis (IPA). After subsetting the results of URA to transcriptional regulators, we found that IPA predicted a variety of transcription factors to be differentially activated in either of the technologies, and that these transcriptional changes were coordinated across cell types (Fig. 4d). For example, we found TFs *XBP1, ATF4* and *ATF6,* which are activated upon endoplasmic reticulum stress, and *NFE2L2* and *NRF1,* which mediate the oxidative stress response and are involved in maintaining mitochondria redox homeostasis^25,60,61^ to be upregulated in 10x. These predictions are in line with our previous findings (Fig. 2b, Supplementary Fig. 2a), demonstrating a higher proportion of stressed cells in 10x compared to Parse. Since we found that the Parse dataset had a larger proportion of reads originating from TFs, we decided to extend our analysis to gene regulatory network (GRN) analysis using SCENIC^62^. Average area under the curve (AUC) scores per cell type and technology were z-score normalized and subjected to k-means clustering (Fig. 4e). We found that the two technologies clustered apart (column clusters 1 and 3 for 10x, and 2 and 4 for Parse) but also cell types divided into two meta groups based on the activity of GRNs (column clusters 1 and 2 were enriched in neurons, while column clusters 3 and 4 contained predominantly more progenitor cell types, Fig. 4e). Below, we highlight differences in regulon activity of specific TFs between both technologies and cell types.

For example, cell type-specific regulon activity is found in column clusters 3 and 4 (cell types: RL, PAX6 RL, RL-div, Prog-div, Prog 1, Unknown 1, Unknown 2), and especially dividing RL and progenitor cells. These cells had elevated z-scores for the E2F family, which are TFs involved in cell cycle progression and apoptosis^63^ (Fig. 4e, Supplementary Fig. 4c). In the same row cluster as the E2F family of TFs (row cluster 4), there was regulon for NFIA, a TF involved in GC maturation during cerebellar development^64,65^ (Fig. 4e, Supplementary Fig. 4c). Conversely, column clusters 1 and 2 (cell types: HindExN, Prog 2, GPC, GPC/GC, HindN, DAB1/CALB1/CALB2 HindN, Newborn PC), were enriched for ZEB1, a marker of neuronal migration necessary for the proper development of various brain regions and tumorigenesis in pediatric patients, including medulloblastoma^66^ (Fig. 4e, Supplementary Fig. 4c). Specifically newborn PCs were enriched for GBX2 and LHX5 (Fig. 4e). GBX2 is a known homeobox gene that plays a significant role in cerebellar regionalization^67^, and LHX5 is one of the TFs that define PC cell fate^68^ (Fig. 4e, Supplementary Fig. 4c). Collectively, SCENIC analysis revealed cell type-specific regulon activity characteristic for distinct cerebellar cell types irrespective of the technology used for cell capture. Hence both technologies can be used for GRN inference.

Although cell type-specific regulon activity signatures could be observed in both technologies, there were also regulons with differential activity between technologies (e.g., a subset of regulons in row cluster 2, Fig. 4e, Supplementary Fig. 4c). Examples of such regulon activity signatures were SCAND1 and ZNF580 regulons, two TFs known for their involvement in the cellular response to hypoxic stress^69,70^ but also in mitochondrial and ribosomal functions (Fig. 4e, Supplementary Fig. 4c).

Collectively, with our secondary analysis, we confirmed the previous findings that 10x cells had higher expression of ribosomal and mitochondrial protein-coding genes as defined by quality control compared to Parse cells (i.e., gene name pattern “RPS/RPL” for ribosomal and “MT-” for mitochondrial protein-coding genes). Furthermore, we found that other genes with mitochondrial and ribosomal functions were significantly deregulated in the 10x dataset. Additionally, URA predicted a coordinated change in the activity of cellular stress-related transcriptional regulators between 10x and Parse datasets. These findings suggest that 10x cells have a higher expression of cell stress-related transcriptional signatures, and Gruffi-based exclusion of cells with high stress scores did not solve the problem entirely. Finally, SCENIC analysis revealed that regulons are differentially active between cell types in both technologies. Hence, transcriptional differences between technologies did not mask transcriptional differences between cell types.

## Discussion

In this study, we compared two broadly used and commercialized approaches for sample multiplexing of scRNA-seq: 10x Genomics (10x) and Parse Biosciences (Parse). We generated cerebellar organoids, as an example of a complex 3D tissue that requires dissociation, to comprehensively explore the strengths and limitations of each technology. Regionalized neural organoids, such as cerebellar organoids, are commonly used in neuroscience research but can be challenging due to heterogeneity between samples, batches, and iPSC lines. Therefore they require in-depth characterization, for example, by multiplexed scRNA-seq^11,18^. To compare scRNA-seq datasets across experiments and studies conducted in different labs and to differentiate technical and biological causes of variance, it is essential to understand artefacts and biases introduced by different experimental pipelines of the capture techniques. Specifically, we differentiated three control iPSC lines into cerebellar organoids according to a published protocol^15^. Organoids were pooled and dissociated at D35 and D50, and the cells were split into two aliquots, one of which was subjected to the 10x and the other to the Parse multiplexing and sequencing pipelines. The two methods were then compared regarding library efficiency, differential transcript capture, cell type enrichment, and the information obtained from comprehensive secondary analysis.

Sample preparation between the two technologies differs considerably: while cells are kept alive for a longer time until lysis in the 10x workflow, Parse samples are fixed directly after dissociation. Consequently, Parse samples do not have to be processed in parallel providing more flexibility during sample processing and allowing the handling of higher sample numbers in one sequencing run. Therefore, we suggest that this approach is advantageous for larger experimental designs.

We compared the technical sequencing parameters of both methods. We found that the average cell recovery rate differed considerably between the two techniques. While 42.7% of cells were recovered in the 10x workflow, only 16.5% recovery was achieved in Parse (Supplementary Fig. 1b). For scarce samples a high cell recovery is clearly beneficial to maximize data output. However, we did not observe the lack of certain cell types within the Parse data set, indicating even cell loss across all cell types.

For both methods, most reads were mapped to the genome. However, we observed differences in the number of genes detected and their properties. In accordance with the previous study comparing Parse and 10x on PBMCs^10^, we found that 10x scRNA-seq resulted in a higher number of detected genes, a higher number of protein-coding genes, and a higher number of genes coding for mitochondrial and ribosomal genes compared to Parse (Fig. 1b, Supplementary Fig. 1e). Furthermore, the GC content of captured transcripts was higher in 10x than in Parse. Our analysis also revealed a bias of 10x in capturing shorter transcripts compared to Parse (Fig. 1d, Supplementary Fig. 1h). Moreover, Parse did not only represent longer transcripts but also covered a wider range of gene lengths (Fig. 1b, Supplementary Fig. 1e). Previous functional analysis showed a connection between the transcript length and specific cellular processes and tissue types^71^. While short transcripts are more often associated with skin development and the immune system, longer transcripts more frequently play a role in neuronal development^71^. There is growing evidence for long neural genes to be involved in disease mechanisms during development: long genes are more prone to recurrent double-strand break clusters and are implicated in tumor suppression and psychiatric disorders^72^. Further, long genes can contain broad enhancer-like domains, and their transcription is particularly sensitive to alternations in ASD-associated chromatin regulators. Interestingly, BCL11b (CTIP2) (102,911 bps), a TF crucial for neuronal maturation and differentiation is predicted to be upregulated in Parse in DAB1/CALB1/CALB2 HindN in our data (Fig. 4d). The clinical features of BCL11b-associated neurodevelopmental disorders include ASD, intellectual disability, and cerebellar hypoplasia^74^, which have been previously modeled in organoids^19,20^. These findings indicate that transcript length is a critical technical and biological factor that should be considered when planning scRNA-seq experiments and that Parse could be favorable to investigate differences in long transcripts upon experimental manipulation.

Further, Parse covered a higher number of transcripts encoding TFs among protein-coding genes (Fig. 1b, Supplementary Fig. 1e). To investigate if this bias had effects on GRN we performed GRN analysis SCENIC. Interestingly, Parse generally had higher z-scores for regulons related to neurodevelopment and maturation (Fig. 4e) in contrast to the upregulation of neuron processes assembly-related terms in 10x in GSEA (Fig. 4c). Additionally, we identified regulons that were differentially regulated between cell types and techniques such as *NFIA* which had higher z-scores in RL-derivates in Parse (Fig. 4e) and is involved in GC maturation but also associated with severe neurodevelopmental disorders and gliomas^64,65^. Taken together, the GRN analysis reveals not only cell type but also technique-driven regulon activity. This highlights that identical biological samples result in different analysis results downstream depending on the capture technology.

Regionalized neural organoids have been reported to show high expression of stress pathway-related transcripts due to *in vitro* culturing conditions and insufficient oxygen supply^23,50,51,75^. Additionally, tissue dissociation for single-cell sample preparations is known to induce stress response in dissociated cells^76^.

During QC, we found that the percentage of mitochondrial and ribosomal protein-coding genes was higher in the 10x than in the Parse samples (Fig. 1b), corroborating previous findings^10^. While differences for mitochondrial protein-coding transcripts were minor (10x 3.1% vs Parse 1.7%), the differences for ribosomal protein-coding genes were much more pronounced (10x 17.6% vs Parse 0.5%). The DGE analysis revealed the upregulation of mitochondrial protein-coding genes, and other genes involved in mitochondrial function (Fig. 4b). Hence, the differences in mitochondrial transcripts might be partially explained by higher cell stress in the 10x data and mitochondrial involvement in stress response pathways^77^, rather than having solely technical causes.

To investigate cell stress in cerebellar organoids in more detail, we analyzed the expression of stress-specific modulators. We identified three stress-related modules (oxidative stress, glycolysis, and integrated stress response (ISR)) that separated the two technologies in hierarchical clustering with both technologies showing a stronger module expression at the later time point and 10x demonstrating a higher overall expression of stress modules. It has previously been described that stress-related pathways are enriched in organoids. Cell-intrinsic mechanisms as well as extrinsic factors such as hypoxia can activate the ISR to restore cellular homeostasis. Different cell stressors can also interact with each other to induce the ISR. For example, upon disruption of endoplasmic reticulum (ER) homeostasis, ER stress is induced and can increase the production of reactive oxygen species (ROS) in mitochondria, which induces oxidative stress^78^. These effects can increase during organoid culture as the tissue grows, which may explain the elevated stress response-associated transcriptional signature at D50 compared to D35 of differentiation (Fig. 2b). Since stressed cells are frequently found in scRNA-seq datasets of organoids, a powerful bioinformatic approach called Gruffi was developed to remove cells with a high cell stress signature from neural organoid datasets^50^. Applying Gruffi to our dataset revealed a noticeably higher percentage of stressed cells in the 10x compared to the Parse dataset at both time points (Fig. 2d). This might stem from the difference in the handling of dissociated cells in the two technologies. In the Parse procedure, cells are fixed directly after dissociation, thus limiting the induction of the expression of stress genes. In contrast, live cells are undergoing the 10x capture, prolonging the period between dissociation and cell lysis during capture, which might increase stress-related responses of live cells found in 10x data. Interestingly, this effect is more pronounced in D50 than in D35 samples indicating that more mature neural cells are more susceptible to the mechanical stress of dissociation and live processing in 10x. These findings suggest that identical samples of cerebellar organoids show a technology and time point-specific stress response reflected by striking differences in the number of cells identified as stressed cells by the Gruffi algorithm (Fig. 2d). Further, we found DEGs, especially mitochondrial and ribosomal protein-coding transcripts between the two technologies across all clusters and gene set enrichment analysis in GPCs revealed deregulation of GO terms related to nucleotide processing and mitochondrial respiration (Fig. 4c). To explore which upstream mechanisms could have led to these transcriptional changes, we performed URA. Interestingly, URA for transcriptional regulators predicted the upregulation of ER-stress pathways related TFs *XBP1*, *ATF4*, and *ATF6* as well as oxidative stress mediators *NFE2L2* and *NRF1* in 10x compared to Parse^78^. Together these results suggest that not only the hypoxic culture conditions of organoids^51^ but also the single-cell dissociation and capturing pipeline may induce cell stress. The cell capture technology used thus affects the output data obtained from biologically identical samples, and this effect should be considered when interpreting and comparing organoid data to reference datasets.

To investigate the biological reproducibility of the organoid differentiation protocol, we assessed the percentage of cells identified as neural based on reference-query mapping with human developmental transcriptome^52^. This analysis showed a commitment towards neural fate in 52.1% of all cells, suggesting the initial tissue specification could be improved in the differentiation protocol. Different neural organoid protocols^14,79^ and a recently published protocol for cerebellar organoids^29^ use dual SMAD inhibition during the initiation of differentiation to prevent meso– and endodermal fates thus promoting neural induction^80^. In contrast, the cerebellar differentiation protocol used in this study employs only one SMAD pathway inhibitor, the TGFß-inhibitor SB-4321542^15^. Dual SMAD inhibition might enhance neuroectodermal commitment in cerebellar organoids.

To date, studies employing transcriptional analysis of cerebellar organoids have used only one iPSC line^27,29^. Interestingly, we observed noticeable differences between the differentiation efficiency of the three control cell lines, with the KOLF2.1J-derived cerebellar organoids demonstrating the lowest number of neural cells, especially pronounced at D50. Considering that all three cell lines were differentiated in parallel to minimize technical confounder effects, this finding implicates that iPSC line-inherent mechanisms can influence the differentiation efficiency^81^. This finding highlights the need to use isogenic control iPSCs when analyzing pathogenic variants^82^. Addressing the heterogeneous outcomes of differentiation protocols, a recent study suggests adjusting concentrations of small molecules and growth factors in a cell line-specific manner to decrease the proportion of mesodermal off-target tissue for the differentiation of cortical organoids^25^. A similar approach could potentially alleviate differences in neuroectodermal fate commitment during cerebellar differentiation across the three iPSC lines used in this study. Taken together, new protocols should be tested and optimized with multiple control iPSC lines to ensure robustness of differentiation efficiency^83^. Despite the differences between the three iPSC lines used in this study, we demonstrated that cerebellar organoids generated cerebellar cells of both RL and VZ lineage. Comparing our data set with a recently published cerebellar organoid transcriptomic dataset^29^ revealed general agreement with our annotation indicating a similar cellular population resulting from different protocols. However, the cerebellum is a complex brain region with various cell types^37^ and to what extend different cerebellar organoid protocols recapitulate the whole cerebellum or rather specific regions like the cerebellar nuclei or cerebellar cortex remains to be investigated.

In conclusion, our comprehensive comparison of Parse and 10x scRNA-seq sample multiplexing and cell capture strategies encompassed library efficiency, differential transcript capture, cell type preferences, and secondary analysis outcomes, showing distinct strengths and limitations of each method. While both methods provide the experimental benefits of sample multiplexing, we revealed significant differences between the two strategies. Overall, our findings indicate that while 10x provided higher cell recovery and gene detection rates, Parse captured longer transcripts and a wider range of transcript lengths and resulted in lower cell stress. Minimizing cell stress is especially relevant in the context of regionalized neural organoids, in which cell stress may be an important artefact^51^. Our detailed secondary analyses demonstrated that these technical differences have relevant biological implications. These insights are crucial for selecting the most suitable scRNA-seq multiplexing technology based on specific research goals. Future studies should consider these factors to improve the accuracy and biological relevance of single-cell transcriptomic analyses. Finally, we demonstrated cerebellar organoid differentiation and in-depth characterization on three iPSC lines and highlighted the importance of employing several cell lines in these studies to encompass cell line-dependent heterogeneity and to produce robust results.

## Methods

### iPSC culture

Commercially available iPSC lines BIONi010-C (Source: EBiSC), BIONi037-A (Source: EBiSC) and KOLF 2.1J (Source: The Jackson Laboratory) were cultured under standard conditions (37°C, 5% CO2, and 100% humidity) in E8 Flex medium (BIONi010-C and BIONi037-A) and mTeSR plus (STEMCELL Technologies, Cat. no 100-0276) (Gibco, Cat. no. A2858501) on hESC-qualified growth factor-reduced Matrigel-coated (Corning, Cat. no. 354277) cell culture dishes (Greiner, Cat. no. 657160). Passaging was performed in colonies using Gentle Dissociation Reagent (STEMCELL Technologies, Cat. no. 07174) once the culture reached 80%-90% confluency. The culture medium was supplemented with Thiazovivin (Sigma-Aldrich, Cat. no. 420220) upon passaging for one day. All cell lines were tested for mycoplasma contamination using PCR Mycoplasma Detection Set (TaKaRa, Cat. no. 6601) and maintained under passage 20. The pluripotency for each cell line was confirmed by immunocytochemistry against OCT4 (rabbit, 1:500, Abcam, Cat. no. ab19857) prior to the start of differentiation.

### Generation of cerebellar organoids

Cerebellar organoids were generated as previously described^84^ with some modifications: 80-90% confluent iPSCs were dissociated into single cells using Accutase (Merck, Cat. no. A6964), and 4,500 cells were seeded per well of 96 well V-bottom low adhesion plates (S-bio, Cat. no. MS-9096VZ) in E8 Flex medium (Gibco, Cat. no. A2858501), supplemented with 10 μM Y-27632 (Cayman Chemical, Cat. no. 10005583). Once the aggregates reached a diameter of 250 μm, the medium was changed to growth factor-free chemically defined medium (gfCDM) supplemented with 50 ng/ml FGF2 (PeproTech, Cat. no. 100-18B) and 10 μM SB-431542 (Tocris, Cat. No. 1614), and this day was considered day 1 of differentiation (D1). At D7, FGF2 and SB-431542 were reduced to 33.3 ng/ml and 6.67 μM, respectively. At D14, media was supplemented with 100 ng/ml FGF19 (PeproTech, Cat. No. 100-32). The medium was changed to Neurobasal Medium at D21, supplemented with 300 ng/ml SDF-1 from D28 to D34. From D35 onwards, media was changed to complete BrainPhys (StemCell Technologies, Cat. no. 5793), supplemented with 10 μg/ml BDNF (PeproTech, Cat. no. 450-02), 100 μg/ml GDNF (PeproTech, Cat. no. 450-10), 100 mg/ml dbcAMP (PeproTech, Cat. no. 1698950) and 250 mM ascorbic acid (Tocris, Cat. no. 4055). All three cell lines were processed in parallel during differentiation, single-cell dissociation, and sequencing.

### Single-cell dissociation of cerebellar organoids, library preparation, and sequencing

On D35 and D50, 24 organoids per cell line were pooled and dissociated using the Papain dissociation kit (Worthington, Cat.No. LK003150) following a published protocol with minor modifications^14^. Cells were counted, and cell suspensions were split into two parts for further processing.

Samples for the 10x Genomics (10x) pipeline were labeled with cell multiplexing oligos (CMO, 10x Genomics, Cat. no. 1000261) according to the manufacturer’s instructions and subsequently pooled at an equal ratio. The cell count for the cell suspension was determined, and the sample was loaded onto two lanes of a Chromium Next Gen Chip G (10x Genomics, Cat. no. 1000120) with a targeted cell recovery of 12,000 (D35) and 14,000 (D50) cells per lane. Library preparation was performed with the Chromium Next GEM Single Cell 3’ kit v3.1 (10x Genomics, Cat. no. 1000268), and sequencing was performed on NovaSeq 6000 with S1 flow cell kit and 100 cycles (Illumina, Cat. no. 20028319).

Samples for Parse Bioscience (Parse) workflow were fixed according to the manufacturer’s instructions using the Evercode fixation kit for cells (Parse Bioscience, Cat. No. WF300). Fixed Parse samples were stored at –80°C until all samples were harvested. The samples were characterized by the day of differentiation (D35 or D50) and cell line (BIONi010-C, BIONi037-A, or KOLF2.1J). Every sample was loaded as a technical duplicate into 2 independent wells, with all samples spanning wells 1-12. Sequencing was performed using a molarity of 62.4 nM and 3% PhiX spike in on the Nova Seq 6000 with SP flow cell kit and 200 cycles (Illumina).

### Data downsampling, preprocessing, and quality control

Initially, the datasets from 10x and Parse pipelines had different sequencing depths and cell numbers (Supplementary Table 1). To compare the two technologies fairly, we downsampled datasets from both technologies to an average of 50,000 reads per cell. The FASTQ files were downsampled with the *seqtk sample* tool, and the same seed was applied for the forward and reverse reads. For Parse data, FASTQ files from each of the 2 sub-libraries were demultiplexed into 6 samples. Using *split-pipe* (v1.1.2), the samples were preprocessed, aligned, sorted, annotated, and passed to a DGE (here, digital gene expression), resulting in a count matrix. Afterwards, the 2 sub-libraries were merged with the corresponding *combine* mode of *split-pipe*. For 10x data, read downsampling was performed for individual libraries. Afterwards, downsampled FASTQ files were processed with *cellranger* (v.7.2.0) *multi* pipeline, and cells were assigned with their cell line of origin based on their CMO.

Gene names in gene expression matrices between the two technologies were harmonized in the following manner. Firstly, ENSEMBL gene identifiers were used to merge expression matrices. Secondly, ENSEMBL identifiers were replaced by HGCN identifiers wherever possible (41,980 genes), and ENSEMBL identifiers were used in other cases (20,930 genes). The merged gene expression matrix was further converted into Seurat objects (Seurat v.5.1.0). Gene biotypes were retrieved from bioMart using ENSEMBL gene identifiers. Ribosomal protein-coding genes were identified using HGCN gene names starting from RPS/RPL. Mitochondrial protein-coding genes were identified using HGCN gene names starting from MT-. The percentage of gene expression for ribosomal and mitochondrial protein-coding genes as well as for individual gene biotypes were calculated using *PercentageFeatureSet()*. For calculating the percentage of counts originating from transcription factors (TF) among protein-coding genes, the count matrix was first subset to protein-coding genes, and *PercentageFeatureSet()* was applied to this matrix using the list of human TFs^85^.

Next, quality control (QC) was performed on cell and gene levels. Cells were excluded if one of the following criteria was met: (1) number of individual genes per cell ≤ 2,000; (2) number of individual genes per cell ≥ 13,000; (3) number of genes per UMI ≤ 0.8; and (4) percentage of mitochondrial genes ≥ 8%. We excluded genes from the expression matrices when their cumulative expression across all cells was ≤ 8. No ambient RNA and doublet removal were performed.

### Data normalization, clustering, integration, and dimensionality reduction

After QC, data were normalized using *NormalizeData()* function from Seurat with default parameters. Normalized data were scaled, and principal component analysis (PCA) was performed based on the z-scaled expression of the 2,000 most variable features (*FindVariableFeatures()*). Additionally, normalized counts were integrated using *IntegrateData()* function with reciprocal PCA (RPCA). Dimensionality reduction and clustering were performed using both un– and integrated data. *RunUMAP()* function was used to perform dimensionality reduction with 30 neighbors and 30 principal components (PC). Louvain clustering was performed using *FindClusters()* function.

### Technology-specific analyses: correlation analysis, transcript length, and GC content

To analyze the correlation of gene expression between technologies, we used cells that passed quality control, averaged the expression of genes for each technology, and calculated Pearson’s correlation coefficient. Differentially expressed genes (DEG) between technologies were identified using MAST algorithm implemented in *FindMarkers()* function as previously described^10^ with the following cutoffs: absolute log2 fold change (log2FC) > 1, adjusted p-value < 0.01. Gene length and GC content were retrieved from bioMart.

### Cellular stress assessment

Normalized unintegrated counts were used to analyze the expression of cell stress-related gene ontology (GO) terms using *AddModuleScore()* function. We also generated a random set of genes of mean GO term size and analyzed the expression of these genes as a module to use as an internal control for module expression analysis. Hierarchical clustering was performed on mean module expression of cell stress-related GO terms across samples.

Gruffi cell stress analysis was performed using normalized unintegrated counts following the authors’ instructions^50^. Firstly, two GO terms were chosen for negative selection: glycolytic process (GO:0006096) and integrated stress response signaling (GO:0140467); and one for positive selection: neurogenesis (GO:0022008). Next, module expression of the selected GO terms was analyzed in “granules”, and 90% quantile threshold was chosen for selecting stressed cells.

### Germ layer assessment

Normalized integrated counts were used to perform Azimuth reference-query mapping^53^ of our dataset with human fetal development transcriptome^52^. Cells were further classified as “neural” and “non-neural” based on cell type assignment from Azimuth (Supplementary Table 3).

Gruffi differentiation lineage analysis was performed using normalized integrated counts. Firstly, two GO terms were chosen for negative selection: endoderm (GO:0001706) and mesoderm (GO:0001707) formation; and two for positive selection: nervous system development (GO:0007399) and neurogenesis (GO:0022008). Next, module expression of the selected GO terms was analyzed in “granules”, and 90% quantile threshold was chosen for selecting neural and non-neural cells.

### Neural data processing and cell type annotation

After germ layer assessment, the dataset was subset to neural cells by labels originating from Azimuth reference-query mapping and further downsampled to retain the equal number of cells in 10x and Parse datasets (7,212 cells per technology). Data normalization, clustering, integration, and dimensionality reduction workflow steps were repeated as described above.

VoxHunt^57^ was used to analyze the brain region identity of the cells. 10 genes with the highest area under the curve (AUC) scores per brain region of the developing mouse brain at E15 were retrieved, resulting in 186 unique regional marker genes. These marker genes were used to assess the similarity of gene expression profiles between our samples and BrainSpan human developmental transcriptome^58^ at postconceptional weeks 12 and 13.

Cell type annotation was performed for clusters at resolution 0.9 by a combination of approaches: (1) retrieving cluster marker genes by *FindAllMarkers()* with MAST (normalized counts provided as input) and ROC (raw counts provided as input) algorithms; (2) visualizing expression of canonical marker genes for cell types in the developing mouse and human cerebellum.

### Reference-query mapping with published primary cerebellar development and cerebellar organoids transcriptomic datasets

For reference-query mapping of our cells that were classified as neural, we first used human cerebellar development transcriptomic dataset^38^ as a reference. We downsampled the reference dataset to 1,000 cells per cell type as defined by the metadata (author_cell_type column). Secondly, we normalized, found variable features, scaled, and performed PCA on both reference and query datasets using Seurat default parameters. Integration was performed using *FindTransferAnchors()* function with the “pcaproject” option and 30 PCs. Predicted cell type annotations and prediction scores were obtained from *TransferData()* function wrapped into *MapQuery()* with default parameters and reference label being “author_cell_type”. Integration with the cerebellar organoids transcriptomic dataset was performed as described above with minor modifications: (1) the complete reference dataset was used for mapping; (2) the reference label was “final.clusters”.

### Differential gene expression analysis and functional enrichment analysis

For differential gene expression (DGE) analysis, the raw counts originating from neural cells were used. First, cells were grouped by cell type, technology, cell line, and day of differentiation, and groups smaller than 20 cells were omitted from further analysis. Gene counts were aggregated by technology, cell line, and day of differentiation using *AggregateExpression()* function with a default option to calculate the sum of raw counts per cell group. Importantly, we did not further downsample our dataset to generate an equal number of cells per cell group. The aggregated counts were used as samples for *DESeq2* (v.1.42.1) differential gene expression analysis between technologies within individual cell types^86^. Log2FC were shrunken using *apeglm* shrinkage estimator^87^ as implemented in DESeq2. Volcano plots were generated using *EnhancedVolcano* library (v.1.20.0).

Gene set enrichment analysis (GSEA) with GO terms was performed by *clusterProfiler* (v.4.10.1)^88^ using biological processes ontology as input, gene set size of 50 to 500 genes, false discovery rate (FDR) as a p-value adjustment method, and the threshold for q-value of 0.05. For significantly deregulated GO terms, similarity matrices were calculated and simplified using the *binary cut* approach implemented in *simplifyEnrichment* (v.1.12.0) package^89^.

### Upstream regulator analysis

Upstream regulator analysis was performed using Ingenuity Pathway Analysis (IPA) software (Qiagen). Briefly, cell type-specific DESeq2 output matrices were used for IPA core analysis with the following cutoffs: (1) absolute log2FC > 1; (2) q-value < 0.0001. For visualizations, molecule type was restricted to transcription regulators, and bias-corrected z-scores across cell types were used for hierarchical clustering using the *ComplexHeatmap* package (v.2.18.0). When z-scores were unavailable, they were assigned to 0.

### Gene regulatory network (GRN) activity analysis

We performed GRN analysis closely following the official pySCENIC protocol^62,90^. First, the annotated raw count matrix produced with Seurat and the list of human TFs were processed, inferencing importance values or the weights of regulatory interactions between TFs and target genes. Second, the inferred interactions (“adjacencies”) were searched in the cisTarget databases to identify the enriched binding motifs. Third, TFs and target genes indicated by the enriched motifs were grouped into regulons (regulatory modules of the network). Finally, the regulons were assessed for the enrichment in each cell. With the count matrix as a source of the expression data, cells were assigned scores, i.e., AUC, of the activity levels of their regulons. Z-scores were further calculated based on AUC scores of individual regulons, and k-means clustering of z-scores was performed to reveal groups of co-regulated regulons. Regulon target genes and GO Biological Processes were used for gene set overrepresentation analysis (ORA) by clusterProfiler (v.4.10.1) with gene set size of 5 to 500 genes, false discovery rate (FDR) as a p-value adjustment method, and the threshold for q-value of 0.1.

### Statistics

R v.4.3.2 was used for statistical analysis. Statistical tests are described in text and figure legends. Two-sided unpaired t-tests were used to compare two groups. For comparisons with more than two groups, we used three-way ANOVA. Within a set of comparisons (e.g., for quality control metrics), the Benjamini-Hochberg method of p-value adjustments was used.

## Figure legends

**Supplementary Fig. 1.**
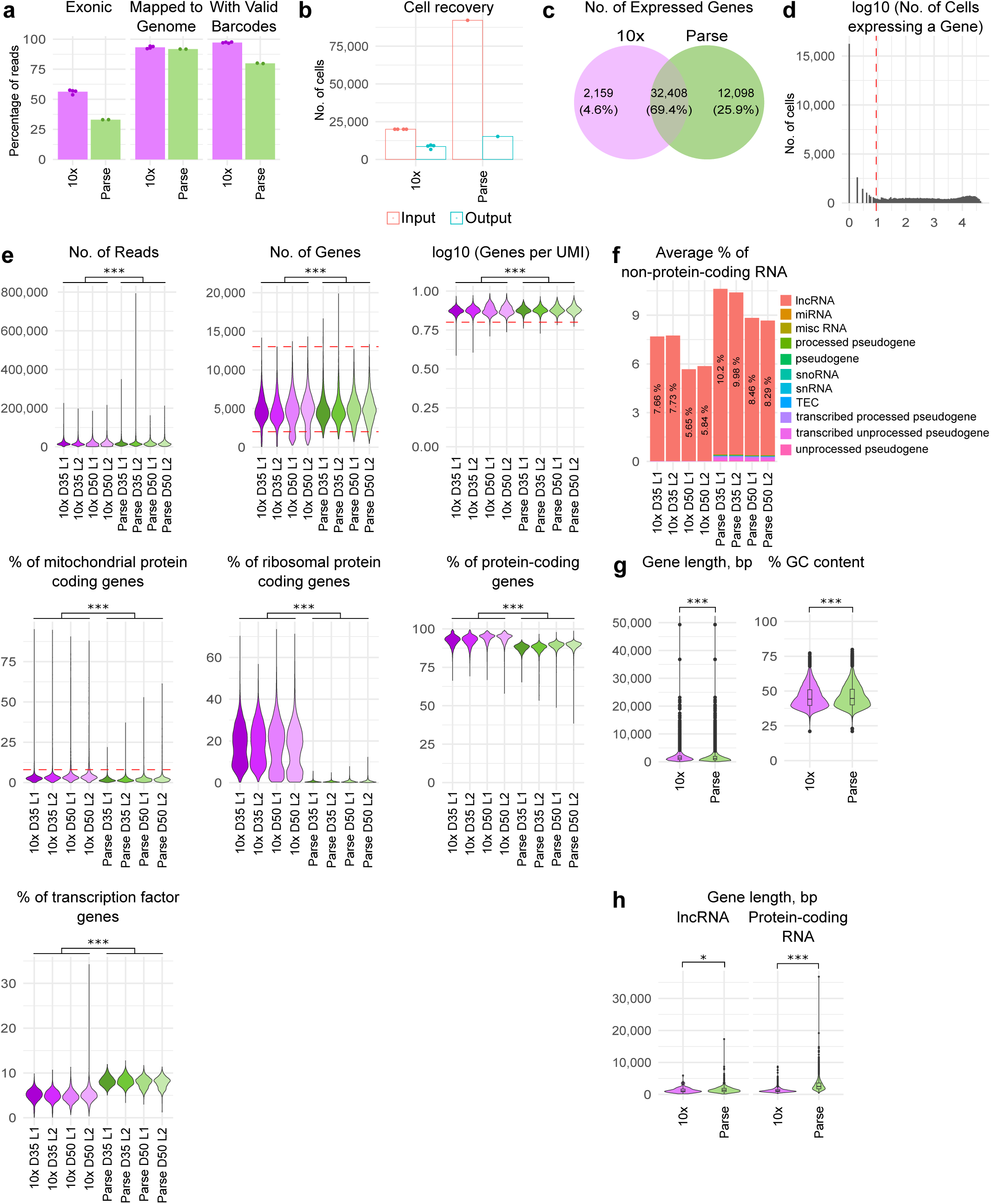
Complementary to Fig. 1. Quality control and gene quantification biases in the data. **a**, Percentage of raw reads mapping to exonic regions, genome, and having valid barcodes. Bars represent the mean; dots represent the individual libraries. **b**, Numbers of input and output cells. Bars represent the mean; for 10x data, dots represent individual libraries; for Parse data, dots represent the total number of cells in the experiment. **c**, Venn diagram of genes expressed in at least 1 cell in each of the two technologies. Color represents technology. **d**, Distribution of the number of cells expressing a gene. **e**, Quality statistics before quality control. Red dashed lines represented threshold values. Color represents sample identity with respect to technology (10x or Parse), day of differentiation (D35 or D50), and library (L1 or L2). 10x, n = 33,951, Parse, n = 15,226 cells. Three-way ANOVA, p-values represent differences between technologies, *** p < 0.001. **f**, Stacked bar plot representing average proportion of reads originating from non-protein-coding RNAs (ncRNA). Color represents ncRNA biotype. **g**, Distributions of gene GC content and gene length for all genes expressed in either of the two technologies. Two-sided t-test, *** p < 0.001. **h**, Distributions of gene length for differentially expressed genes per gene biotype between technologies. Two-sided t-test, * p < 0.05, *** p < 0.001.

**Supplementary Fig. 2.**
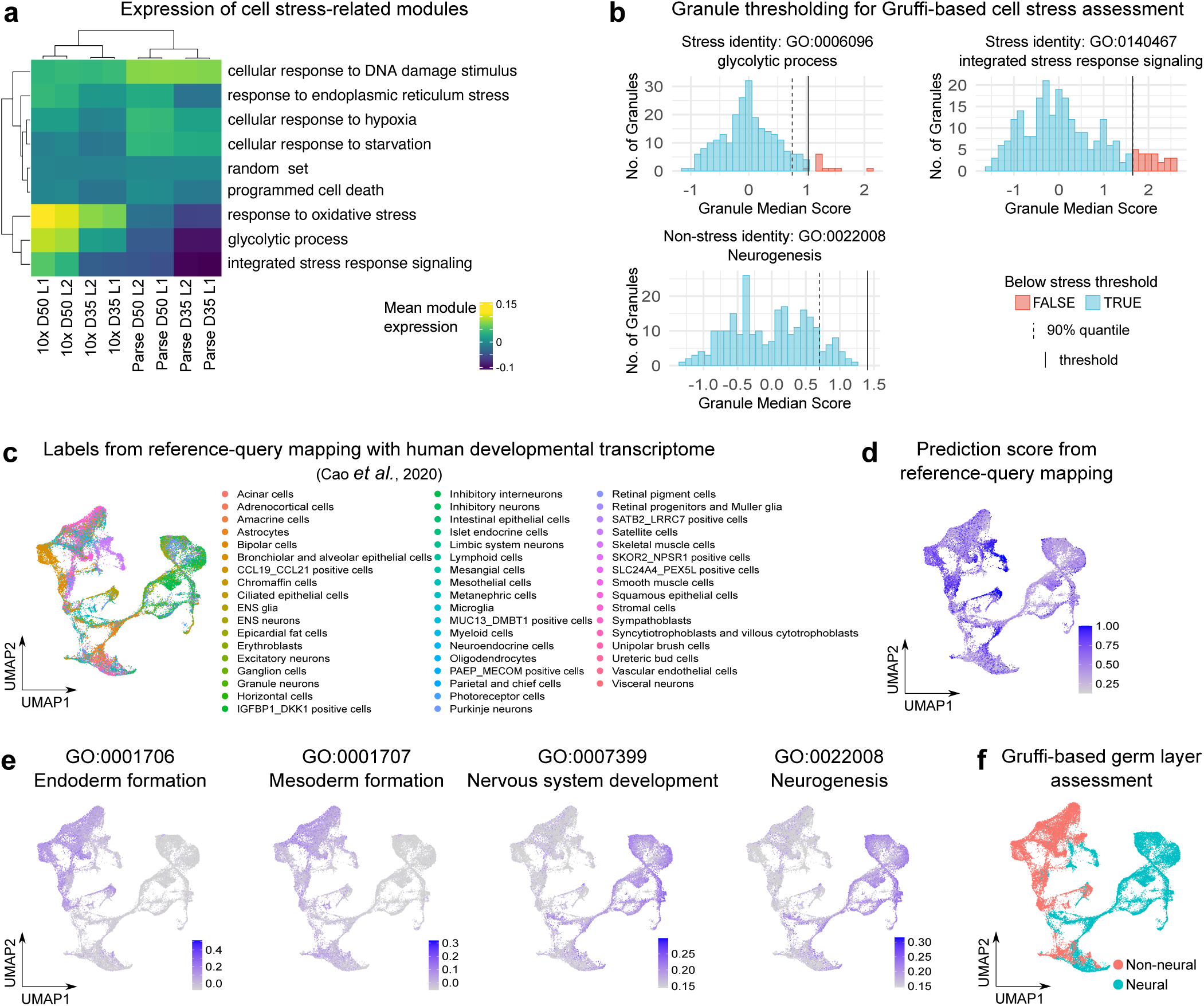
Complementary to Fig. 2. Assessment of neural lineage identity. **a**, Heatmap representing mean module expression scores of gene ontology terms related to aspects of cell stress. **b**, Histograms representing distribution of granule scores for expression of cell stress modules. Color represents stress classification. Solid black line represents stress threshold. Dashed black line represents 90% quantile of the distribution of granule expression score. Dashed blue and red lines represent median values of non– and stressed cells. **c**, UMAP plot representing cell type identity as assigned based on reference-query integration with human developmental transcriptome^52^. **d**, Feature plot showing prediction score based on reference-query integration with human developmental transcriptome. **e**, Feature plots showing module expression scores for GO terms guiding Gruffi-based lineage identity assessment. **f**, UMAP plot representing neural lineage status of cells based on Gruffi-based lineage identity assessment. Three-way ANOVA, p-values represent differences between technologies, n.s. p ≥ 0.05, * p < 0.05, ** p < 0.01, *** p < 0.001.

**Supplementary Fig. 3.**
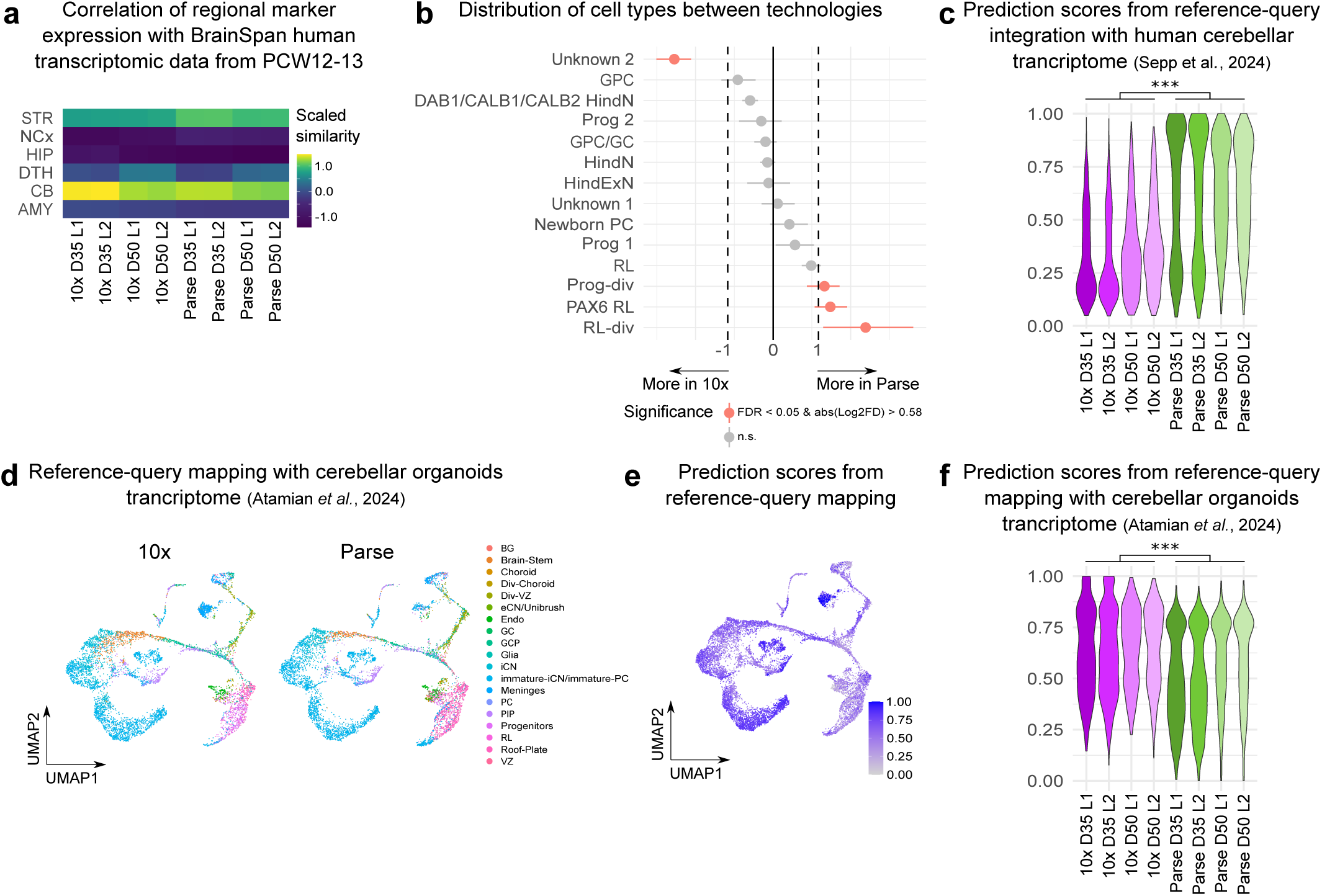
Complementary to Fig. 3. Assessment of regional identity and cell type annotation. **a**, Heatmap of scaled similarity metric of VoxHunt algorithm comparing samples with human neocortical RNA-seq data from BrainSpan using brain regional markers obtained from Mouse Brain Atlas at E13. **b**, Permutation test on cell type composition of cerebellar organoids. Differentially abundant cell types are represented in pink. Cell types with FDR less than 0.05 and absolute log2 fold change more than 0.58 were considered differentially abundant. **c**, Distribution of prediction scores based on reference-query integration with human cerebellar transcriptome^38^. **d**, UMAP plot representing cell type identity as assigned based on reference-query integration with cerebellar organoids transcriptome^29^. **e**, Feature plots showing prediction score based on reference-query integration with cerebellar organoids transcriptome^29^. **f**, Distribution of prediction scores based on reference-query integration with human cerebellar organoids. For **c** and **f**, color represents sample identity with respect to technology (10x or Parse), day of differentiation (D35 or D50), and library (L1 or L2). For **c** and **f** three-way ANOVA, p-values represent differences between technologies, *** p < 0.001.

**Supplementary Fig. 4.**
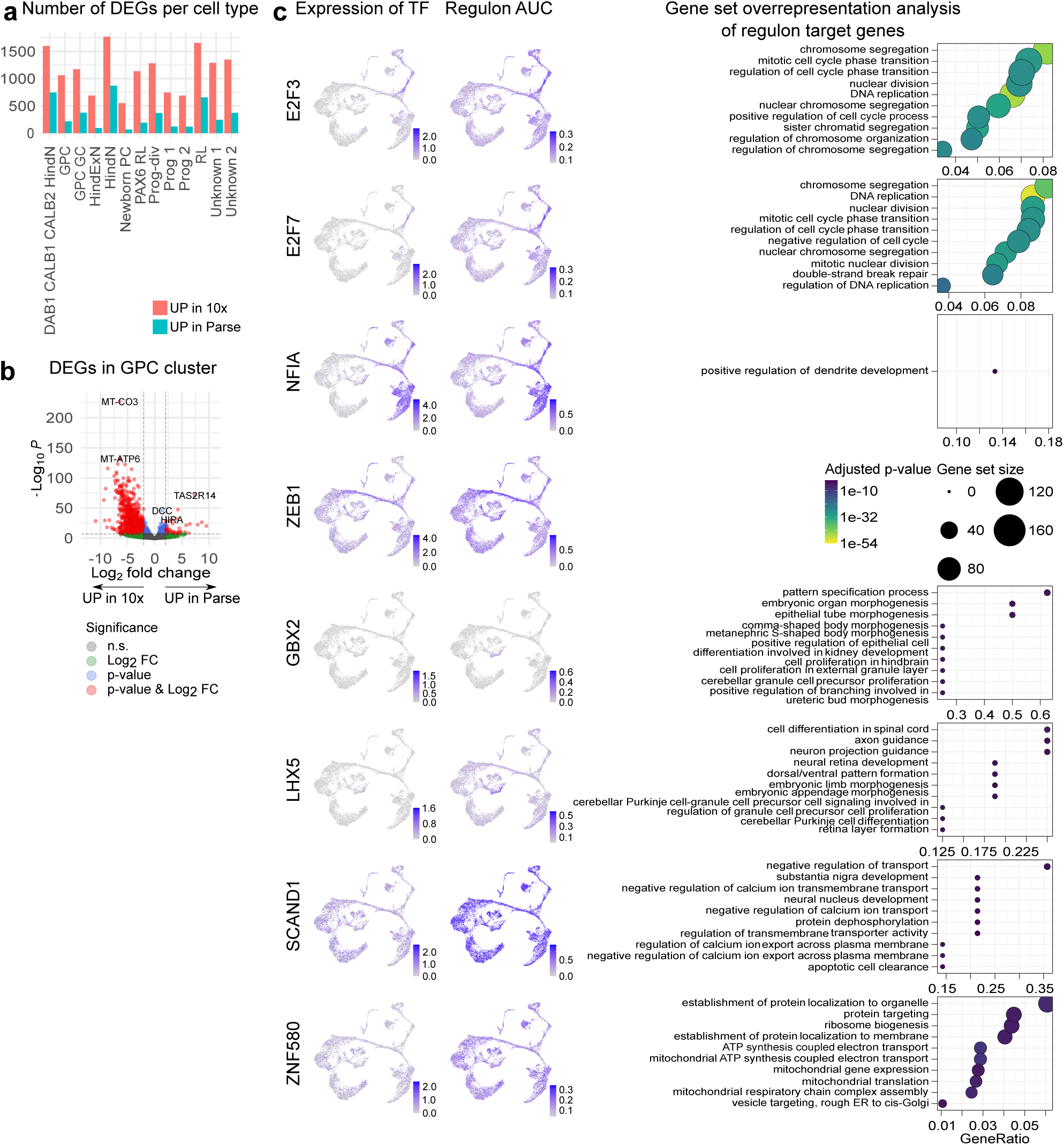
Complementary to Fig. 4. Differential gene expression between technologies. **a**, Bar plot representing number of differentially expressed genes per cell type. **b**, Volcano plot representing differential gene expression in GPC cluster without genes that are exclusively expressed in one of the technologies. **c**, Feature plots showing expression of selected TFs (left column), their regulon AUC scores (middle column), and results of gene set overrepresentation analysis in TF target genes within regulons (right column). ZEB1 did not have any significantly enriched GO terms.

## Data and code availability

Code and data will be made available upon peer-reviewed publication of the manuscript.

## Authors’ contributions

**KS**: Conceptualization, Methodology, Software, Formal analysis, Writing – Original draft, Writing – Review & Editing, Visualization, Project administration; **TK**: Conceptualization, Methodology, Investigation, Writing – Original draft, Writing – Review & Editing, Visualization, Project administration; **VL**: Methodology, Software, Formal analysis, Writing – Original draft; Writing – Review & Editing; **ZY**: Investigation, Writing – Original draft; Writing – Review & Editing; **KB**: Investigation; Writing – Review & Editing; **JM**: Funding acquisition; Writing – Review & Editing; **NC**: Conceptualization, Methodology, Formal analysis, Writing – Original draft, Writing – Review & Editing, Resources, Supervision; **SM**: Conceptualization, Methodology, Writing – Review & Editing, Resources, Supervision, Funding acquisition.

## Acknowledgements

We thank Antje Schulze-Selting, Elisabeth Gustafsson, Christina Kulka, and Ezgi Atay for technical support. We thank Christopher Sifuentes, Yogesh Singh, and Vincent Hammer for strategic and technical discussions.

We are grateful for financial support from the Hertie Foundation (Gemeinnützige Hertie-Stiftung), the Ministerium für Wissenschaft, Forschung und Kunst Baden-Württemberg state postgraduate fellowship (to TK), Add-on Fellowship of the Joachim Herz Foundation (to KS), and the Heidelberger Akademie der Wissenschaften (WIN Kolleg). NGS sequencing methods were performed with the support of the DFG-funded NGS Competence Center Tübingen (INST 37/1049-1). This project has been made possible in part by grant number 2022-316727 from the Chan Zuckerberg Initiative DAF, an advised fund of Silicon Valley Community Foundation. This research has been partially funded by the Deutsche Forschungsgemeinschaft (DFG, German Research Foundation) under Germany’s Excellence Strategy via the Excellence Cluster 3D Matter Made to Order (EXC-2082/1 – 390761711).

## References

1. Li, C. et al. Single-cell brain organoid screening identifies developmental defects in autism. Nature 621, 373–380 (2023).

2. Van de Sande, B. et al. Applications of single-cell RNA sequencing in drug discovery and development. Nat Rev Drug Discov 22, 496–520 (2023).

3. Camp, J.G., Wollny, D. & Treutlein, B. Single-cell genomics to guide human stem cell and tissue engineering. Nat Methods 15, 661–667 (2018).

4. Evrony, G.D., Hinch, A.G. & Luo, C. Applications of Single-Cell DNA Sequencing. Annu Rev Genomics Hum Genet 22, 171–197 (2021).

5. Yasen, A. et al. Progress and applications of single-cell sequencing techniques. Infect Genet Evol 80, 104198 (2020).

6. Luecken, M.D. & Theis, F.J. Current best practices in single-cell RNA-seq analysis: a tutorial. Mol Syst Biol 15, e8746 (2019).

7. Wolock, S.L., Lopez, R. & Klein, A.M. Scrublet: Computational Identification of Cell Doublets in Single-Cell Transcriptomic Data. Cell Syst 8, 281–291 e9 (2019).

8. Rosenberg, A.B. et al. Single-cell profiling of the developing mouse brain and spinal cord with split-pool barcoding. Science 360, 176–182 (2018).

9. Cheng, J., Liao, J., Shao, X., Lu, X. & Fan, X. Multiplexing Methods for Simultaneous Large-Scale Transcriptomic Profiling of Samples at Single-Cell Resolution. Adv Sci (Weinh) 8, e2101229 (2021).

10. Xie, Y. et al. Comparative Analysis of Single-Cell RNA Sequencing Methods with and without Sample Multiplexing. International Journal of Molecular Sciences 25, 3828 (2024).

11. Camp, J.G. & Treutlein, B. Human organomics: a fresh approach to understanding human development using single-cell transcriptomics. Development 144, 1584–1587 (2017).

12. Tang, X.Y. et al. Human organoids in basic research and clinical applications. Signal Transduct Target Ther 7, 168 (2022).

13. Lancaster, M.A. & Knoblich, J.A. Generation of cerebral organoids from human pluripotent stem cells. Nat Protoc 9, 2329–40 (2014).

14. Velasco, S. et al. Individual brain organoids reproducibly form cell diversity of the human cerebral cortex. Nature 570, 523–527 (2019).

15. Silva, T.P. et al. Scalable Generation of Mature Cerebellar Organoids from Human Pluripotent Stem Cells and Characterization by Immunostaining. J Vis Exp (2020).

16. Susaimanickam, P.J., Kiral, F.R. & Park, I.H. Region Specific Brain Organoids to Study Neurodevelopmental Disorders. Int J Stem Cells 15, 26–40 (2022).

17. Renner, H. et al. A fully automated high-throughput workflow for 3D-based chemical screening in human midbrain organoids. Elife 9(2020).

18. Eichmüller, O.L. & Knoblich, J.A. Human cerebral organoids — a new tool for clinical neurology research. Nature Reviews Neurology 18, 661–680 (2022).

19. Sarieva, K. et al. Human brain organoid model of maternal immune activation identifies radial glia cells as selectively vulnerable. Mol Psychiatry 28, 5077–5089 (2023).

20. Kagermeier, T. et al. Human organoid model of pontocerebellar hypoplasia 2a recapitulates brain region-specific size differences. Disease Models & Mechanisms 17(2024).

21. Giorgi, C. et al. Brain Organoids: A Game-Changer for Drug Testing. Pharmaceutics 16(2024).

22. Corsini, N.S. & Knoblich, J.A. Human organoids: New strategies and methods for analyzing human development and disease. Cell 185, 2756–2769 (2022).

23. Qian, X., Song, H. & Ming, G.L. Brain organoids: advances, applications and challenges. Development 146(2019).

24. Andrews, M.G. & Kriegstein, A.R. Challenges of Organoid Research. Annu Rev Neurosci 45, 23–39 (2022).

25. Bertucci, T. et al. Improved Protocol for Reproducible Human Cortical Organoids Reveals Early Alterations in Metabolism with MAPT Mutations. bioRxiv (2023).

26. Silva, T.P. et al. Transcriptome profiling of human pluripotent stem cell-derived cerebellar organoids reveals faster commitment under dynamic conditions. Biotechnology and Bioengineering 118, 2781–2803 (2021).

27. Nayler, S., Agarwal, D., Curion, F., Bowden, R. & Becker, E.B.E. High-resolution transcriptional landscape of xeno-free human induced pluripotent stem cell-derived cerebellar organoids. Sci Rep 11, 12959 (2021).

28. Muguruma, K. Self-Organized Cerebellar Tissue from Human Pluripotent Stem Cells and Disease Modeling with Patient-Derived iPSCs. Cerebellum 17, 37–41 (2018).

29. Atamian, A. et al. Human cerebellar organoids with functional Purkinje cells. Cell Stem Cell 31, 39–51 e6 (2024).

30. Schmahmann, J.D. The cerebellum and cognition. Neurosci Lett 688, 62–75 (2019).

31. Zhang, P. et al. The cerebellum and cognitive neural networks. Front Hum Neurosci 17, 1197459 (2023).

32. Sathyanesan, A. et al. Emerging connections between cerebellar development, behaviour and complex brain disorders. Nat Rev Neurosci 20, 298–313 (2019).

33. Mapelli, L., Soda, T., D’Angelo, E. & Prestori, F. The Cerebellar Involvement in Autism Spectrum Disorders: From the Social Brain to Mouse Models. Int J Mol Sci 23(2022).

34. (!!! INVALID CITATION !!!).

35. Harada, H., Sato, T. & Nakamura, H. Fgf8 signaling for development of the midbrain and hindbrain. Dev Growth Differ 58, 437–45 (2016).

36. Lowenstein, E.D., Cui, K. & Hernandez-Miranda, L.R. Regulation of early cerebellar development. FEBS J 290, 2786–2804 (2023).

37. Leto, K. et al. Consensus Paper: Cerebellar Development. The Cerebellum 15, 789–828 (2016).

38. Sepp, M. et al. Cellular development and evolution of the mammalian cerebellum. Nature 625, 788–796 (2024).

39. Haldipur, P. et al. Spatiotemporal expansion of primary progenitor zones in the developing human cerebellum. Science 366, 454–460 (2019).

40. van der Heijden, M.E. & Sillitoe, R.V. Cerebellar dysfunction in rodent models with dystonia, tremor, and ataxia. Dystonia 2(2023).

41. Kamei, T. et al. Survival and process outgrowth of human iPSC-derived cells expressing Purkinje cell markers in a mouse model for spinocerebellar degenerative disease. Experimental Neurology, 114511 (2023).

42. Coolen, M. et al. Recessive PRDM13 mutations cause fatal perinatal brainstem dysfunction with cerebellar hypoplasia and disrupt Purkinje cell differentiation. Am J Hum Genet 109, 909–927 (2022).

43. Kresbach, C. et al. Intraventricular SHH inhibition proves efficient in SHH medulloblastoma mouse model and prevents systemic side effects. Neuro Oncol 26, 609–622 (2024).

44. van Essen, M.J. et al. PTCH1-mutant human cerebellar organoids exhibit altered neural development and recapitulate early medulloblastoma tumorigenesis. Dis Model Mech 17(2024).

45. Ballabio, C. et al. Modeling medulloblastoma in vivo and with human cerebellar organoids. Nat Commun 11, 583 (2020).

46. Muguruma, K., Nishiyama, A., Kawakami, H., Hashimoto, K. & Sasai, Y. Self-organization of polarized cerebellar tissue in 3D culture of human pluripotent stem cells. Cell Rep 10, 537–50 (2015).

47. Salmen, F. et al. High-throughput total RNA sequencing in single cells using VASA-seq. Nature Biotechnology 40, 1780–1793 (2022).

48. Zheng, W., Chung, L.M. & Zhao, H. Bias detection and correction in RNA-Sequencing data. BMC Bioinformatics 12, 290 (2011).

49. Tirosh, I. et al. Dissecting the multicellular ecosystem of metastatic melanoma by single-cell RNA-seq. Science 352, 189–196 (2016).

50. Vértesy, Á. et al. Gruffi: an algorithm for computational removal of stressed cells from brain organoid transcriptomic datasets. The EMBO Journal 41, e111118 (2022).

51. Bhaduri, A. et al. Cell stress in cortical organoids impairs molecular subtype specification. Nature 578, 142–148 (2020).

52. Cao, J. et al. A human cell atlas of fetal gene expression. Science 370, eaba7721 (2020).

53. Hao, Y. et al. Integrated analysis of multimodal single-cell data. Cell 184, 3573–3587.e29 (2021).

54. Olson, E.N. MyoD family: a paradigm for development. Genes Dev 4, 1454–1461 (1990).

55. Itoh, N. FGF10: A multifunctional mesenchymal–epithelial signaling growth factor in development, health, and disease. Cytokine & Growth Factor Reviews 28, 63–69 (2016).

56. Wilson, S.W. & Houart, C. Early Steps in the Development of the Forebrain. Developmental Cell 6, 167–181 (2004).

57. Fleck, J.S. et al. Resolving organoid brain region identities by mapping single-cell genomic data to reference atlases. Cell Stem Cell 28, 1148–1159.e8 (2021).

58. Miller, J.A. et al. Transcriptional landscape of the prenatal human brain. Nature 508, 199–206 (2014).

59. Aldinger, K.A. et al. Spatial and cell type transcriptional landscape of human cerebellar development. Nat Neurosci 24, 1163–1175 (2021).

60. Chen, X., Shi, C., He, M., Xiong, S. & Xia, X. Endoplasmic reticulum stress: molecular mechanism and therapeutic targets. Signal Transduct Target Ther 8, 352 (2023).

61. Jager, R., Bertrand, M.J., Gorman, A.M., Vandenabeele, P. & Samali, A. The unfolded protein response at the crossroads of cellular life and death during endoplasmic reticulum stress. Biol Cell 104, 259–70 (2012).

62. Aibar, S. et al. SCENIC: single-cell regulatory network inference and clustering. Nature Methods 14, 1083–1086 (2017).

63. DeGregori, J. The genetics of the E2F family of transcription factors: shared functions and unique roles. Biochimica et Biophysica Acta (BBA) – Reviews on Cancer 1602, 131–150 (2002).

64. Dini, G. et al. NFIA haploinsufficiency: case series and literature review. Front Pediatr 11, 1292654 (2023).

65. Chen, K.S., Lim, J.W.C., Richards, L.J. & Bunt, J. The convergent roles of the nuclear factor I transcription factors in development and cancer. Cancer Lett 410, 124–138 (2017).

66. Fratini, L. et al. Oncogenic functions of ZEB1 in pediatric solid cancers: interplays with microRNAs and long noncoding RNAs. Molecular and Cellular Biochemistry 476, 4107–4116 (2021).

67. Su, C.Y., Kemp, H.A. & Moens, C.B. Cerebellar development in the absence of Gbx function in zebrafish. Dev Biol 386, 181–90 (2014).

68. Zhao, Y. et al. LIM-homeodomain proteins Lhx1 and Lhx5, and their cofactor Ldb1, control Purkinje cell differentiation in the developing cerebellum. Proceedings of the National Academy of Sciences 104, 13182–13186 (2007).

69. Tweedie, D. et al. Mild traumatic brain injury-induced hippocampal gene expressions: The identification of target cellular processes for drug development. Journal of Neuroscience Methods 272, 4–18 (2016).

70. Yin, C. et al. RNA-seq Analysis Reveals Potential Molecular Mechanisms of ZNF580/ZFP580 Promoting Neuronal Survival and Inhibiting Apoptosis after Hypoxic-ischemic Brain damage. Neuroscience 483, 52–65 (2022).

71. Lopes, I., Altab, G., Raina, P. & de Magalhaes, J.P. Gene Size Matters: An Analysis of Gene Length in the Human Genome. Front Genet 12, 559998 (2021).

72. Wei, P.C. et al. Long Neural Genes Harbor Recurrent DNA Break Clusters in Neural Stem/Progenitor Cells. Cell 164, 644–55 (2016).

73. Zhao, Y.T. et al. Long genes linked to autism spectrum disorders harbor broad enhancer-like chromatin domains. Genome Res 28, 933–942 (2018).

74. Seigfried, F.A. & Britsch, S. The Role of Bcl11 Transcription Factors in Neurodevelopmental Disorders. Biology (Basel) 13(2024).

75. Pasca, A.M. et al. Human 3D cellular model of hypoxic brain injury of prematurity. Nat Med 25, 784–791 (2019).

76. Neuschulz, A. et al. A single-cell RNA labeling strategy for measuring stress response upon tissue dissociation. Mol Syst Biol 19, e11147 (2023).

77. Manoli, I. et al. Mitochondria as key components of the stress response. Trends Endocrinol Metab 18, 190–8 (2007).

78. Cao, S.S. & Kaufman, R.J. Endoplasmic reticulum stress and oxidative stress in cell fate decision and human disease. Antioxid Redox Signal 21, 396–413 (2014).

79. Pasca, A.M. et al. Functional cortical neurons and astrocytes from human pluripotent stem cells in 3D culture. Nat Methods 12, 671–8 (2015).

80. Chambers, S.M. et al. Highly efficient neural conversion of human ES and iPS cells by dual inhibition of SMAD signaling. Nature Biotechnology 27, 275–280 (2009).

81. Volpato, V. & Webber, C. Addressing variability in iPSC-derived models of human disease: guidelines to promote reproducibility. Dis Model Mech 13(2020).

82. Pantazis, C.B. et al. A reference human induced pluripotent stem cell line for large-scale collaborative studies. Cell Stem Cell 29, 1685–1702 e22 (2022).

83. Sarieva, K. & Mayer, S. The Effects of Environmental Adversities on Human Neocortical Neurogenesis Modeled in Brain Organoids. Front Mol Biosci 8, 686410 (2021).

84. Silva, T.P. et al. Maturation of Human Pluripotent Stem Cell-Derived Cerebellar Neurons in the Absence of Co-culture. Front Bioeng Biotechnol 8, 70 (2020).

85. Lambert, S.A. et al. The Human Transcription Factors. Cell 172, 650–665 (2018).

86. Love, M.I., Huber, W. & Anders, S. Moderated estimation of fold change and dispersion for RNA-seq data with DESeq2. Genome Biology 15(2014).

87. Zhu, A., Ibrahim, J.G. & Love, M.I. Heavy-tailed prior distributions for sequence count data: removing the noise and preserving large differences. Bioinformatics 35, 2084–2092 (2019).

88. Wu, T. et al. clusterProfiler 4.0: A universal enrichment tool for interpreting omics data. The Innovation 2(2021).

89. Gu, Z. & Hübschmann, D. SimplifyEnrichment: A Bioconductor Package for Clustering and Visualizing Functional Enrichment Results. *Genomics*, Proteomics & Bioinformatics 21, 190–202 (2023).

90. Van De Sande, B. et al. A scalable SCENIC workflow for single-cell gene regulatory network analysis. Nature Protocols 15, 2247–2276 (2020).

